# High-level prediction errors in low-level visual cortex

**DOI:** 10.1101/2023.08.21.554095

**Authors:** David Richter, Tim C Kietzmann, Floris P de Lange

## Abstract

Perception and behaviour are significantly moulded by expectations derived from our prior knowledge. Hierarchical predictive processing theories provide a principled account of the neural mechanisms underpinning these processes, casting perception as a hierarchical inference process. While numerous studies have shown stronger neural activity for surprising inputs, in line with this account, it is unclear what predictions are made across the cortical hierarchy, and therefore what kind of surprise drives this upregulation of activity. Here we leveraged fMRI and visual dissimilarity metrics derived from a deep neural network to arbitrate between two hypotheses: prediction errors may signal a local mismatch between input and expectation at each level of the cortical hierarchy, or prediction errors may incorporate feedback signals and thereby inherit complex tuning properties from higher areas. Our results are in line with this second hypothesis. Prediction errors in both low- and high-level visual cortex primarily scaled with high-level, but not low-level, visual surprise. This scaling with high-level surprise in early visual cortex strongly diverges from feedforward tuning, indicating a shift induced by predictive contexts. Mechanistically, our results suggest that high-level predictions may help constrain perceptual interpretations in earlier areas thereby aiding perceptual inference. Combined, our results elucidate the feature tuning of visual prediction errors and bolster a core hypothesis of hierarchical predictive processing theories, that predictions are relayed top-down to facilitate perception.

## Introduction

Predictive processing (PP) theories promise to provide a principled account of cortical computation (Bastos et al., 2012; Friston, 2005, 2009; Rao & Ballard, 1999). One critical ingredient of PP is the computation of prediction errors, i.e. the mismatch between prediction, usually thought of as a top-down signal, and bottom-up input. Such prediction errors then serve as input to the next level in the cortical hierarchy. The brain minimizes prediction errors by recurrently updating its predictions. This process enables the formation of a coherent, stable, and efficient representation of the world. Despite variations in specific predictive processing implementations (Ali et al., 2022; Bastos et al., 2012; Friston, 2005, 2009; Rao & Ballard, 1999; Spratling, 2017), the core concept of prediction error computation is ubiquitous and supported by many empirical observations. For instance, after visual statistical learning, visual cortex is sensitive to the likelihood of an object’s appearance. In particular, activity throughout the ventral visual stream has been shown to be attenuated to expected compared to unexpected appearances of the same stimuli (Egner et al., 2010; Kaposvari et al., 2018; Kok et al., 2012; Richter & de Lange, 2019; Utzerath et al., 2017). This attenuation, also known as expectation suppression, has been observed across different species and modalities (de Lange et al., 2018; Keller & Mrsic-Flogel, 2018; Walsh et al., 2020) and occurs also when predictions and stimuli are task-irrelevant (Meyer & Olson, 2011; Ramachandran et al., 2016; Richter et al., 2018; Schneider et al., 2018). Combined, expectation suppression has frequently been interpreted in the context of PP as reflecting larger prediction errors for unexpected stimuli, and thus taken as crucial evidence that perception fundamentally relies on prediction (de Lange et al., 2018; Walsh et al., 2020).

If prediction and prediction error computations underlie perceptual inference, as suggested by PP theories, we can stipulate that cortical predictions and the associated prediction error signatures must reflect stimulus features that are represented in the respective cortical area. For example, prediction errors in primary visual cortex (V1) may signal deviations from expectation in terms of simple features such as stimulus orientation, edges and contrasts – i.e., visual features that V1 neurons are tuned to (Hubel & Wiesel, 1962). On the other hand, prediction errors in higher visual areas (HVC), for instance in fusiform gyrus, may reflect more complex high-level visual features, such as object identities, spatial relationships between object parts and more abstract concepts such as faces, commonly represented in those areas (Kanwisher et al., 1997; Kravitz et al., 2013; Kriegeskorte et al., 2008). This account suggests that prediction errors mirror *local* feature tuning, unique to each visual cortical area. While some studies have provided indirect support for the feature specificity of sensory prediction errors by investigating tuning specific modulations (Kok et al., 2012; Richter et al., 2022; Yon et al., 2018), little evidence directly shows which visual feature surprise, if in fact any, is reflected in visual prediction errors.

In contrast to local feature tuning, prediction error tuning may be *inherited top-down*. Top-down inheritance is in line with hierarchical PP, because predictions are proposed to be relayed top-down from higher to lower visual areas (Friston, 2005), and thus lower visual areas may come to reflect tuning properties of higher areas in predictive contexts due to the top-down prediction signals. Empirical support for this notion has been obtained in the macaque face processing system (Schwiedrzik & Freiwald, 2017). Schwiedrzik and Freiwald (2017) showed that early areas in the macaque face processing hierarchy inherit viewpoint invariance from later areas when faces are shown in a predictable context. However, whether a similar principle of top-down prediction error tuning inheritance applies across species, to stimuli outside the domain of faces and viewpoints, and importantly across the visual hierarchy remains unknown.

Given that prediction error computation is a core mechanism of PP it is crucial to characterize what kind of visual surprise is tracked by the visual system. Here we aimed to close this gap by exploring features reflected in the visual surprise response after statistical learning. Specifically, we asked (1) whether prediction errors come to reflect any visual feature tuning in predictive contexts, and if so, whether (2) this tuning is in line with the local visual features or inherited top-down. To do so, we exposed human volunteers to images that were either expected or unexpected in terms of their identity, given a preceding cue, while recording whole-brain fMRI. To quantify visual feature surprise across multiple levels of description we used representational dissimilarity metrics derived from a visual deep neural network (DNN). Our results demonstrate that neural responses across multiple visual cortical areas monotonically increased with how visually dissimilar a surprising object was compared to the expected object. Crucially, high-level visual dissimilarity accounted for the surprise induced increase of neural responses, including in the earliest visual cortical area, V1. Prediction errors thus appear to reflect surprise primarily in terms of high-level visual features, demonstrating that earlier visual areas inherit feature tuning usually associated with higher visual areas in predictive contexts, presumably due to feedback signals.

## Results

Human volunteers (n = 33) viewed images that could contain either animate or inanimate entities. Each image was preceded by a letter cue that probabilistically predicted the identity of the image. The expected image was seven times more likely to follow its associated letter cue compared to each of the seven unexpected images. Participants were tasked to classify the content of the image as animate or inanimate. Further details are in the *Methods: Stimuli and experimental paradigm* section.

### Behavioral facilitation by valid prediction

Participants were faster and more accurate when categorizing expected compared to unexpected images (Figure 1). Response times to expected images were faster by an average of 16 ms (Wilcoxon signed rank test: *W*_(32)_ = 15, *z* = -4.74, *p* < 0.001, *r* = -0.947) and more accurate by 1.1% (Wilcoxon signed rank test: *W*_(32)_ = 425.5, *z* = 2.59, *p* = 0.010, *r* = 0.517). This behavioural facilitation thus demonstrated that participants learned and used the underlying statistical regularities to predict inputs. Moreover, accuracy in general was very high (> 95%), indicating effective task compliance.

**Figure 1.**
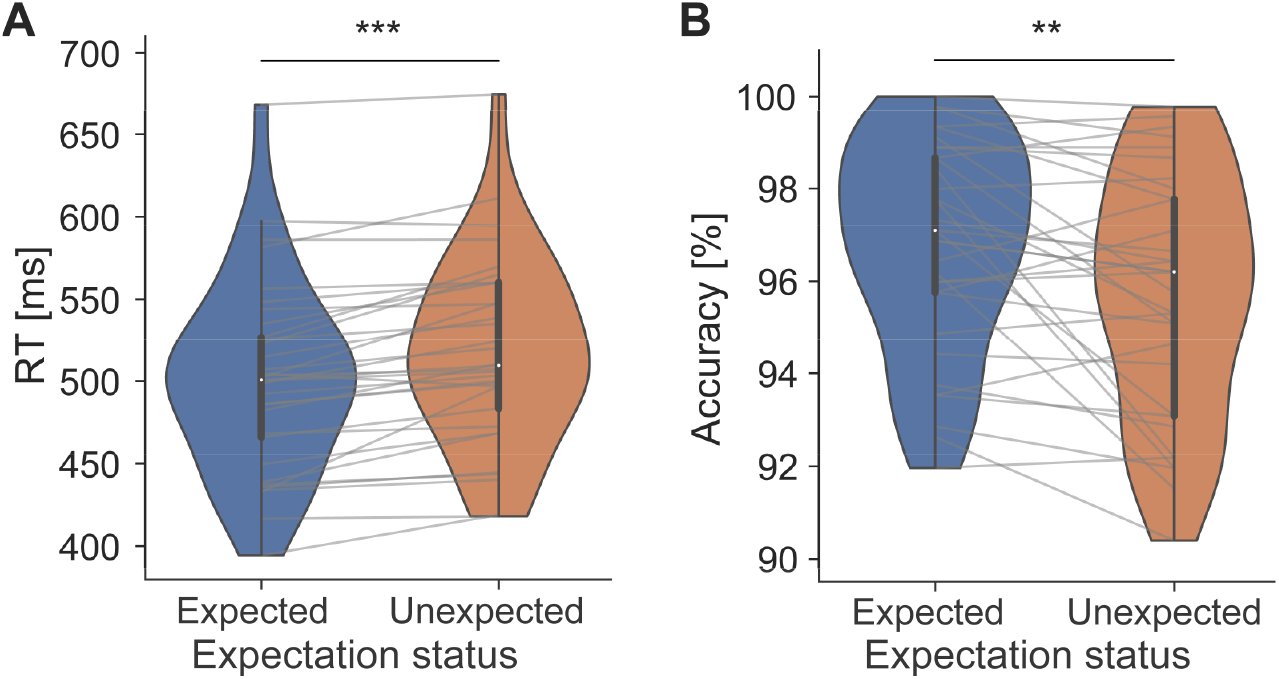
Behavioural facilitation due to prediction. **A)** Reaction times (RT) to expected objects were faster compared to unexpected images. **B)** Accuracy of responses was high overall, but responses were more accurate to expected compared to unexpected images. *** p < 0.001, ** p < 0.01.

### Deep neural network models mirror a gradient from low-to-high level visual features in visual cortex

While (visual) DNNs have been used to successfully explain a variety of neural data (Doerig et al., 2023), we first ensured that the specific feature models utilized here were able to explain visual responses in our data in a prediction-free context. Specifically, we extracted the correlation distance between images from an implementation of AlexNet trained on ecoset (Mehrer et al., 2021). Then we performed representational similarity analysis (RSA) using the representational distances derived from the DNN layers and from the fMRI data obtained during localizer runs. During these runs each image was presented in isolation, without preceding cue and without predictive associations between the images. To assess possible contributions of all DNN layers we repeated this analysis using representational dissimilarity matrices (RDMs) from each of the DNN layers separately. Finally, for each voxel (sphere searchlight) we determined the best DNN layer for explaining neural variance based on the RSA results (correlation coefficient).

Results (Figure 2), showed a gradient from early to higher visual cortex with the corresponding early to late DNN layers best explaining neural variance. Specifically, early visual cortex (EVC) responses were best explained by early layers of the DNN (mostly layers 2-3), intermediate visual areas such as lateral occipital complex (LOC) by intermediate layers (mostly layers 4-6), and HVC responses, for example in the fusiform gyrus, were best explained by late layers of the DNN (layers 7-8). These outcomes affirm previous reports (Guclu & van Gerven, 2015; Mehrer et al., 2021) and validate that our visual feature models, including the low-level (layer 2) and high-level visual model (layer 8), accounted for cortical variance elicited by visual stimulation in an expected pattern of a low-to-high level gradient.

**Figure 2.**
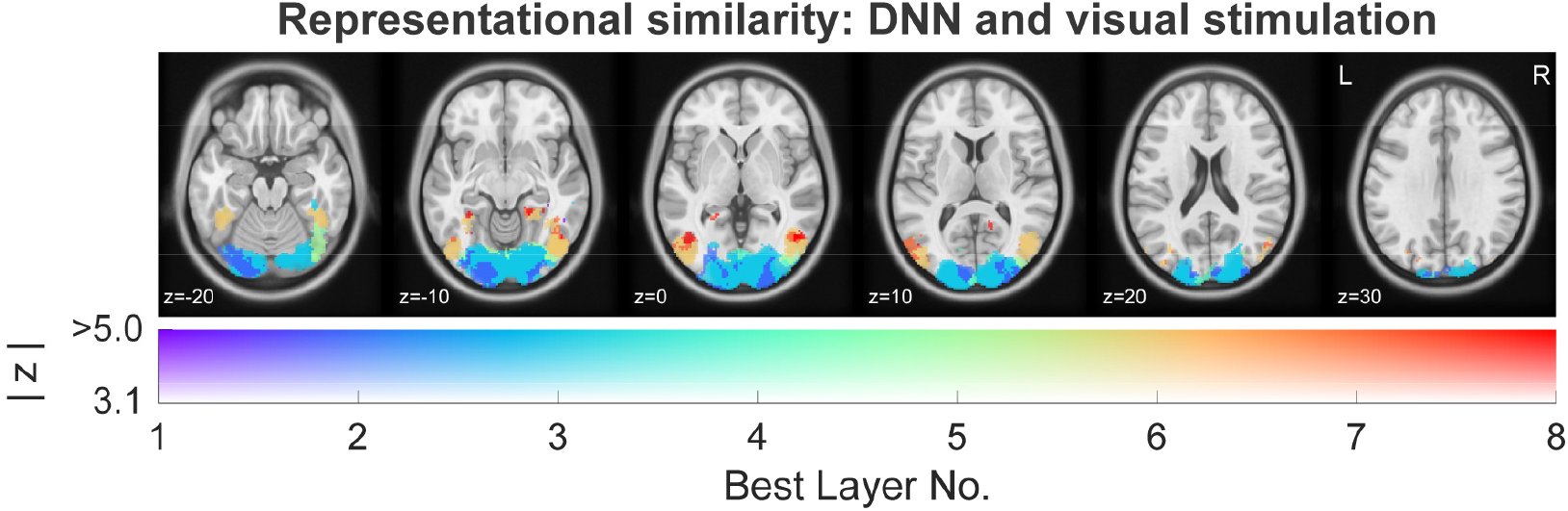
Representational similarity analysis of visual responses in prediction-free contexts. Results show that (feedforward) visual responses during the prediction-free localizer were best explained by a gradient of low-level to high-level visual features going up the ventral visual hierarchy. Cold colors (purple – blue) represent early DNN layers (i.e., low-level visual features), which dominated neural responses in EVC. Warm colors (yellow – red) indicate late DNN layers (i.e., high-level visual features), which best accounted for neural responses in HVC. Analysis was masked to visual cortex and thresholded at z > 3.1 (*p* < 0.001, uncorrected) of the RSA.

### Prediction errors scale with high-level, but not low-level visual feature dissimilarity

Next, we turned our attention to the nature of prediction errors, asking which visual features, if any, they reflect across multiple regions along the ventral visual hierarchy. Figure 3 illustrates the analysis rationale. If low-level visual features, such as local oriented edges and spatial frequency, are predicted in a specific cortical area (e.g., V1) then prediction error magnitudes should scale with low-level surprise in that area. As an example, expecting a specific image of a guitar but seeing an image of another guitar from a different angle should yield a large prediction error as the two images are different from a low-level visual feature standpoint. On the other hand, if high-level visual features, such as more abstract and general guitar features (e.g., a neck and guitar body) are predicted, invariant to local orientation, then the unexpected guitar should not yield a strong prediction error, whereas seeing an image of an unexpected category should result in a large prediction error (even if low-level features are similar). Based on our analyses of DNN alignment with fMRI localizer data and prior work (Mehrer et al., 2021), we decided to use layer 2 of the visual DNN as low-level feature model, and layer 8 (before softmax of the output layer) as high-level visual feature models. This allowed for maximal differentiation of the low-vs high-level models. To model how the BOLD response changed as a function of how visually surprising an unexpected stimulus was in terms of low-level (layer 2) and high-level (layer 8) visual features respectively, we used the dissimilarity of each surprising image, compared to the stimulus expected on that trial, as parametric modulators in the fMRI GLM analysis (for details see *Methods: Data analysis*). In addition, we included multiple control variables, such as task-relevant stimulus animacy and word-level (semantic) dissimilarity.

**Figure 3.**
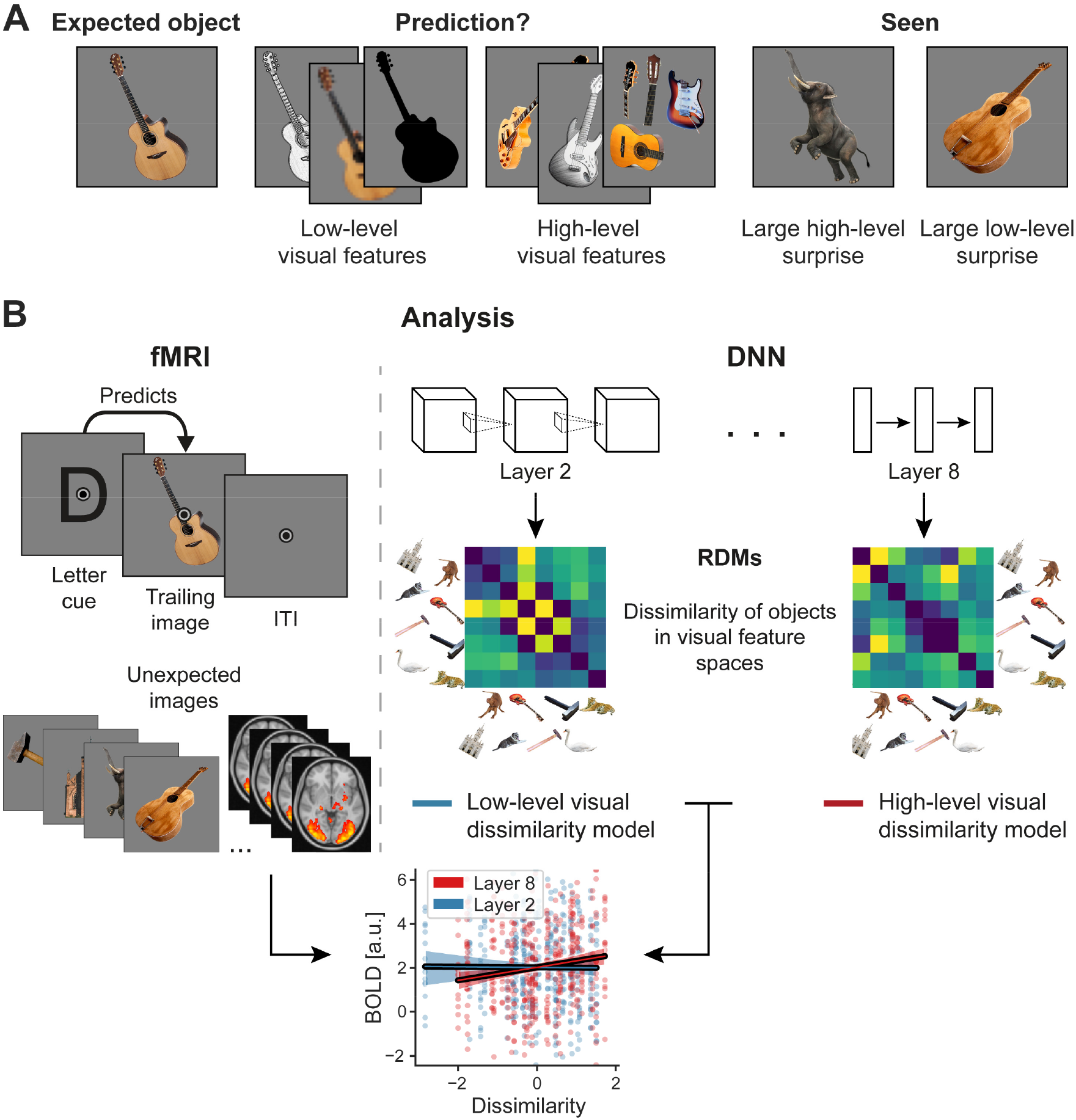
Analytical approach. **A)** If you expect to see the first guitar on the left, what kind of visual features does visual cortex predict? Low-level visual features, illustrated next to the expected image, concern local oriented edges, spatial frequency, and similar properties. High-level visual features entail more abstract visual representations, core object parts and their relationships, such as features shared by all instances of an object, irrespective of the specific depiction. Depending on which features are predicted, the two ‘seen’ images will result in different prediction error magnitudes. The image of the elephant is very different in high-level visual features, but shares some local orientation with the expected guitar, hence resulting primarily in high-level visual surprise. On the other hand, the image of the other guitar is very different in terms of low-level visual features, as it is differently rotated compared to the expected guitar, but it is still a guitar and thus shares high-level visual features. The key question of the analysis is whether and where in the visual system low-level or high-level surprise results in larger prediction errors. **B)** Analysis procedure. The left side shows a single trial with a letter cue predicting a specific image. Below multiple unexpected images are illustrated, which were presented on other trials. The right side depicts the extraction of low-level and high-level visual feature dissimilarity from layer 2 (low-level) and layer 8 (high-level) of the visual DNN. Dissimilarity of the unexpected seen image compared to the expected stimulus was then added as a parametric modulator in the first level GLMs of the fMRI analysis. The graph at the bottom uses data from an example participant to illustrate how BOLD responses in V1 (ordinate) are modulated as a function of low-level (blue) and high-level (red) visual dissimilarity (abscissa). A positive slope thus indicates that the more dissimilar a seen image was relative to the expected stimulus the more vigorous the neural response. In the example data, the image of the unexpected elephant would hence result in larger prediction errors compared to the image of the unexpected guitar, because of the larger high-level surprise. Additional control variables for task relevance (animacy) and word meaning, discussed in more detail later, were also included. We performed this parametric modulation analysis in a voxel-wise fashion across the whole brain.

Results, depicted in Figure 4A, demonstrated that surprise responses scaled significantly with high-level visual dissimilarity (layer 8) in visual cortex, encompassing early and intermediate visual areas (cluster size 779 voxel, 6232 mm^3^; Supplementary Table 1 contains additional details). That is, the more an unexpected stimulus diverged from the expected image in terms of high-level visual features, the more the sensory response increased in magnitude. Surprisingly, we did not find any modulation of neural responses by low-level visual dissimilarity (layer 2) anywhere in visual cortex. In other words, even in EVC prediction error magnitudes were modulated by high-level but not low-level visual surprise, as indexed by layer 2. In the example in Figure 3A this corresponds to the unexpected image of the elephant eliciting a larger prediction error in V1 compared to the unexpected guitar. On the other hand, the low-level surprise elicited by the unexpected guitar would not result in an additional upregulation of prediction errors in visual cortex beyond the associated high-level visual surprise.

**Figure 4.**
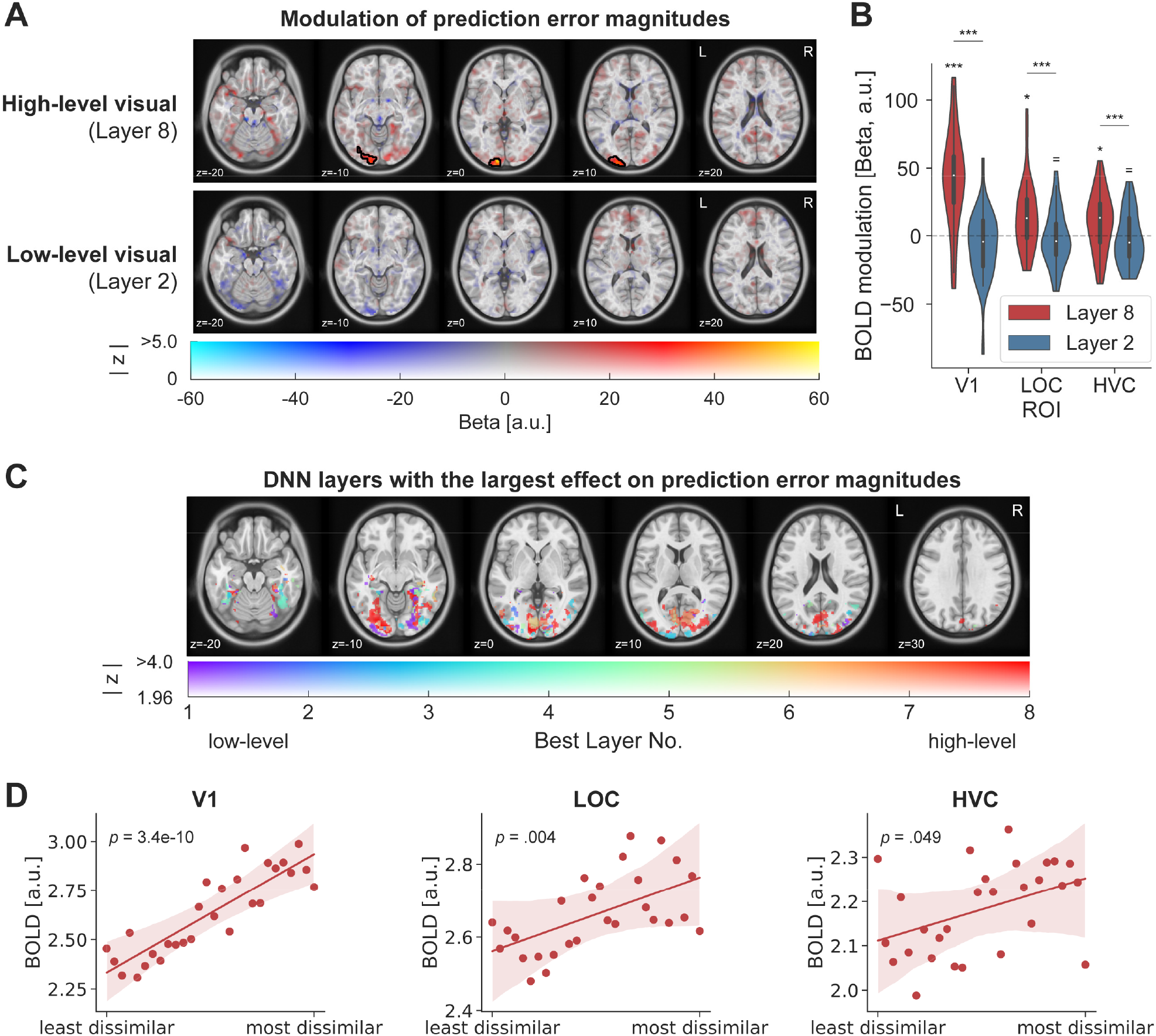
Prediction error magnitude scales with high-level visual feature surprise. **A)** Whole-brain results assessing the modulation of surprise responses as a function of high-level (top row) and low-level (bottom row) visual feature dissimilarity. The top row shows that surprise responses to unexpected images were increased if the image was more distant from the expected image in terms of high-level visual features. Colour indicates the beta parameter estimate of the parametric modulation, with red and yellow representing increased responses. Black outlines denote statistically significant clusters (GRF cluster corrected). No significant modulation of sensory responses was observed by low-level visual surprise. **B)** ROI analysis zooming in on ROIs in early visual (V1), intermediate (LOC) and higher visual cortex (HVC; encompassing occipito-temporal sulcus and fusiform cortex). Results mirror those of the whole-brain analysis, with significant modulations of the visual responses by high-level visual surprise, but not low-level visual surprise. P values are FDR corrected. *** p < 0.001, * p < 0.05, ^=^ BF_10_ < 1/3. **C)** Prediction errors preferentially scale with high-level visual features (layer 8 and layer 7) throughout most of the visual system, including EVC, LOC and HVC. Color indicates the DNN layer with the largest effect (explained variance) on scaling the neural responses to surprising inputs. Cold colors (purple – blue) represent early layers (i.e., low-level visual features), while warm colors (yellow – red) indicate late layers (i.e., high-level visual features). Analysis was masked to visual cortex and thesholded at a liberal z ≥ 1.96 (i.e., *p* < 0.05, two-sided) to explore the landscape of prediction error modulations across DNN layers. Results strongly contrast with those observed for prediction-free visual responses during the localizer (Figure 2). **D)** ROI analysis regressing BOLD responses onto high-level visual dissimilarity. Results show a monotonic relationship between BOLD responses and high-level surprise across all three ROIs. For display purposes dissimilarities were ranked, while statistical inference was performed on the correlation distances.

Our whole-brain results were corroborated by an ROI analysis, depicted in Figure 4B. Results showed a strong difference between high- and low-level visual feature models in modulating BOLD surprise responses (main effect of model: *F*_(1,32)_ = 22.70, *p* < 0.001, 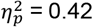). We found reliable modulations of surprise responses by high-level visual, but no significant modulation by low-level visual features, in primary visual cortex (V1: Layer 8: *t*_(32)_ = 6.79, *p* < 0.001, *d* = 1.18; Layer 2: *W* = 190, *p* = 0.159, *d* = -0.32, BF_10_ = 0.58), intermediate visual areas in the lateral occipital complex (LOC: Layer 8: *W* = 126, *p* = 0.017, *d* = 0.55; Layer 2: *t*_(32)_ = -0.68, *p* = 0.504, *d* = -0.12, BF_10_ = 0.23) and high-level visual cortex (HVC: Layer 8: *t*_(32)_ = 2.59, *p* = 0.029, *d* = 0.45; Layer 2: *t*_(32)_ = -0.70, *p* = 0.586, *d* = -0.12, BF_10_ = 0.23). Contrasting the modulation by layer 8 against layer 2 dissimilarity confirmed that layer 8 modulated visual surprise responses significantly more than layer 2 in V1 (*t*_(32)_ = 8.05, *p* < 0.001, *d* = 1.40), LOC (*t*_(32)_ = 4.24, *p* = 0.003, *d* = 0.74), and HVC (*t*_(32)_ = 2.85, *p* = 0.008, *d* = 0.50). Hence, the larger the visual dissimilarity of a surprising stimulus in terms of high-level visual features, the more vigorous the visual response across the ventral visual stream, including V1. No corresponding modulation by low-level visual features was observed, suggesting that high-level features are predominantly reflected in visual prediction error signals.

In a subsequent analysis, we asked which DNN layer explained most neural variance of the prediction error response. We analyzed how neural responses to unexpected stimuli were scaled as a function of surprise indexed by each layer of the DNN. To this end we regressed layer 1-8 dissimilarity onto single trial parameter estimates and determined for each voxel which layer had the largest explained variance. Results (Figure 4C) showed that prediction error magnitudes primarily scaled with high-level visual surprise (layer 8 and layer 7) across most parts of the ventral visual stream, including EVC, LOC, and HVC. We note additional minor clusters in HVC scaled by intermediate layer 4, as well as layer 1 and 3 in EVC and LOC, suggesting that some neural populations scaled with intermediate and low-level surprise as well. In sum, these results present a stark contrast to the modulation of responses during the prediction-free localizer (Figure 2) where a clear gradient from early-to-late layers was observed, with early layers dominating EVC representations. In contrast, prediction errors appear to be scaled preferentially by high-level visual surprise across the visual system, including EVC.

Finally, we assessed the shape of the response modulation by high-level visual surprise by regressing BOLD responses for unexpected stimuli onto layer 8 dissimilarity using an ROI approach. Results, shown in Figure 4D, show that the increased BOLD response to surprising stimuli follows a positive monotonic association across the dissimilarity spectrum in all three ROIs (V1: *t*_(32)_ = 9.35, *p* < 0.001, *d* = 1.63; LOC: *t*_(32)_ = 3.27, *p* = 0.004, *d* = 0.57; HVC: *t*_(32)_ = 2.05, *p* = 0.049, *d* = 0.36). In sum, visual responses across major parts of the ventral visual system appear to monotonically scale with high-level visual surprise.

### Prediction error scaling with high-level visual surprise is not explained by task-relevance, semantic surprise, or inherent properties of the DNN architecture

Multiple alternative explanations could account for a correlation of prediction error magnitudes with high-level visual features. To rule out alternative accounts for our observations, we included multiple control variables as parametric modulators in our GLM analysis. First, layer 8 representations from an untrained (i.e., *random*) but otherwise identical DNN were included. This contrast ruled out that the inherent structure of the DNN architecture or correlations in the input images caused the scaling of prediction errors with late layer representations. Second, *animacy category* was included in the GLM to assess whether high-level visual modulations result from a correlation of high-level visual features with the task-relevant dimension of animacy. Since participants had to distinguish between animate and inanimate entities in the images, prediction error modulations could potentially reflect task responses. Finally, because high-level visual features may significantly correlate with semantic, *word category* level surprise, we also contrasted the high-level visual model against a word2vec (Mikolov, Chen, et al., 2013; Mikolov, Sutskever, et al., 2013) derived model aimed at indexing non-visual, semantic surprise.

These controls revealed that high-level visual surprise (layer 8) best accounted for the data, significantly outperforming an untrained random layer 8 model, animacy category, word category (semantic), and the low-level visual surprise models in explaining prediction error magnitudes (Figure 5A). Statistically significant clusters were found in EVC as well as intermediate visual areas in LOC, and HVC in some contrasts. The exact extent of the modulation varies slightly between contrasts, but overall corroborate that prediction error magnitudes mainly result from high-level visual feature surprise and that none of the control variables likely account for the observed results. Corresponding whole-brain figures contrasting the control parametric modulators against baseline (no modulation) can be found in Supplementary Figure 1.

**Figure 5.**
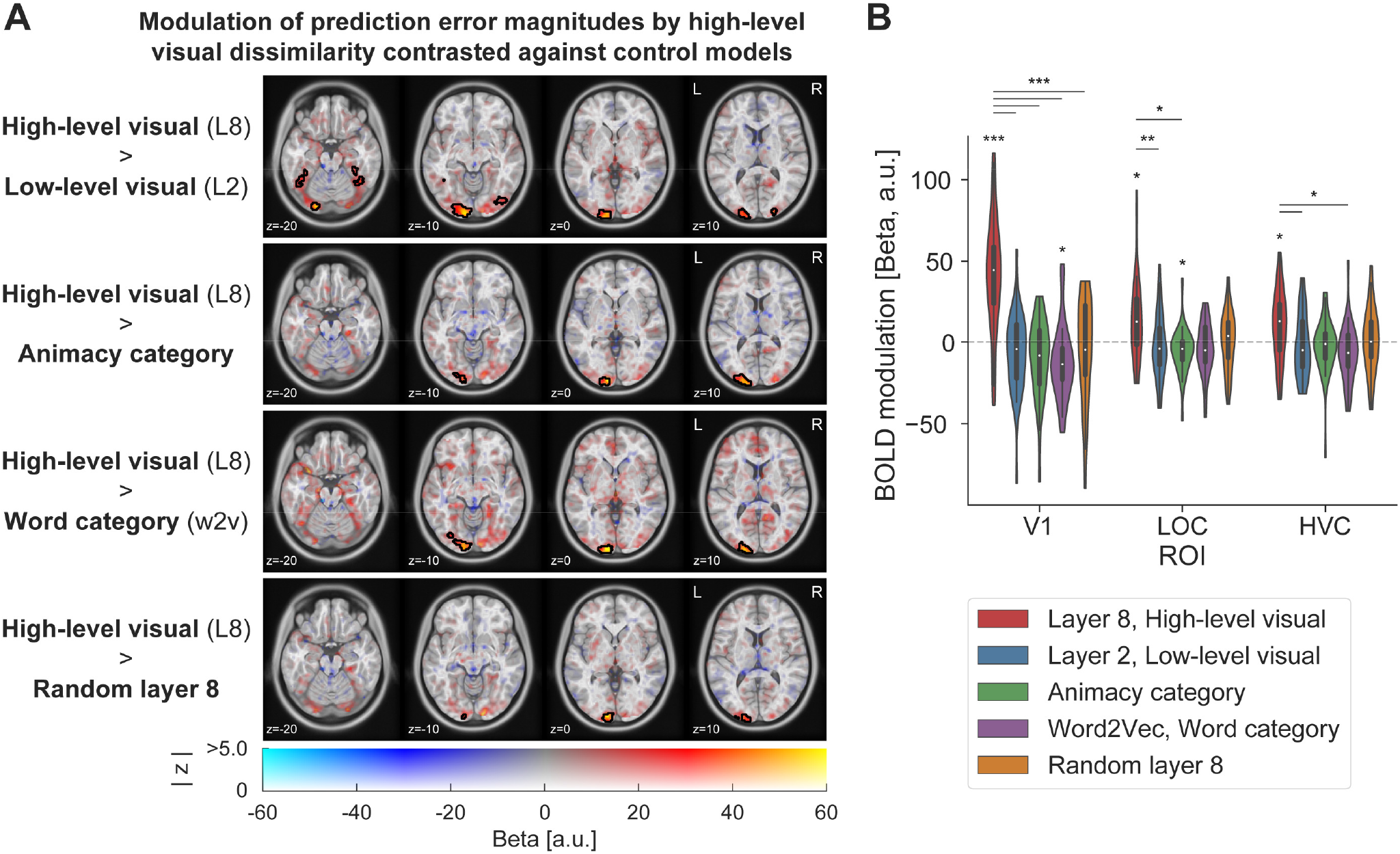
Prediction error magnitudes are best explained by high-level visual feature dissimilarity. **A)** Whole-brain contrasts of the high-level visual feature model (layer 8) contrasted against four control variables. The top row shows that high-level visual models performed significantly better than low-level visual models (layer 2). Similarly, high-level visual surprise better accounted for prediction error magnitudes than the task-relevant animacy category of the unexpected stimuli (second row) and the semantic, word category surprise model (word2vec; third row). The bottom row shows that high-level visual dissimilarity significantly better explained prediction error magnitudes compared to an untrained but otherwise identical DNN layer 8. **B)** ROI analysis including primary (V1), intermediate (LOC) and high-level visual cortex (HVC). Results confirm the whole-brain results, showing significant modulations of BOLD responses by high-level visual surprise compared to low-level visual, response category, and word category surprise. P values are FDR corrected. *** *p* < 0.001, ** *p* < 0.01, * *p* < 0.05.

An ROI analysis (Figure 5B) of the same four contrasts confirmed the whole-brain results. We observed reliable differences between models, which differed across ROIs (main effect of model: *F*_(4,128)_ = 12.20, *p* < 0.001, 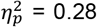; interaction ROI by model: *F*_(4.40,140.93)_ = 11.09, *p* < 0.001, 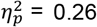). Specifically, we found significantly stronger modulations of BOLD responses by high-level visual dissimilarity compared to all other four parametric modulators in V1 (paired t-tests: all FDR corrected *p* < 0.001; all *d* > 0.99; see Supplementary Table 2 and Supplementary Table 3 for details). Similar, albeit less pronounced results were found in LOC (Layer 8 vs. Layer 2: *p* = 0.001, *d* = 0.74; Layer 8 vs. Animacy category: *p* = 0.010, *d* = 0.61; Layer 8 vs. Word2Vec: *p* = 0.060, *d* = 0.47; Layer 8 vs. Random layer 8: *p* = 0.078, *d* = 0.44) and HVC (Layer 8 vs. Layer 2: *p* = 0.028, *d* = 0.50; Layer 8 vs. Animacy category: *p* = 0.081, *d* = 0.40; Layer 8 vs. Word2Vec: *p* = 0.016, *d* = 0.54; Layer 8 vs. Random layer 8: *p* = 0.189, *d* = 0.32). An additional negative modulation of prediction error magnitudes by word category surprise was observed in V1 (*p* = 0.020, *d* = -0.60), and by animacy category surprise in LOC (*p* = 0.039, *d* = -0.51), suggesting that prediction errors may be attenuated for more semantically dissimilar surprising images in V1 and for stimuli of a different animacy category in LOC compared to the expected image. Finally, to ensure that our results were not dependent on the exact ROI mask size, we repeated the analysis across multiple mask sizes. Results were largely consistent across ROI sizes; see Supplementary Figure 2.

### High-level visual surprise correlates with behavioral response slowing

Participants were asked to classify the contents of the images as animate or inanimate. High-level visual features often correlate with this task-relevant animacy axis. Hence, we assessed whether high-level visual feature surprise might also correlate with increased response times by regressing image dissimilarity onto reaction times to unexpected images. Our findings, shown in Figure 6A, demonstrate that reaction times were indeed modulated by high-level visual (layer 8) dissimilarity. Responses were slower the more dissimilar the surprising images were in high-level visual (*t*_(32)_ = 3.47, *p* = 0.001, *d* = 0.61), but not in low-level visual features (*t*_(32)_ = -1.85, *p* = 0.074, *d* = -0.32; Layer 8 > Layer 2: *t*_(32)_ = 3.10, *p* = 0.004, *d* = 0.54) compared to the expected stimulus. Indeed, the relationship of layer 8 dissimilarity and response slowing was monotonic (Figure 6B). Thus, high-level visual dissimilarity did not only scale with prediction error magnitudes in visual cortex but was also associated with slower RTs.

**Figure 6.**
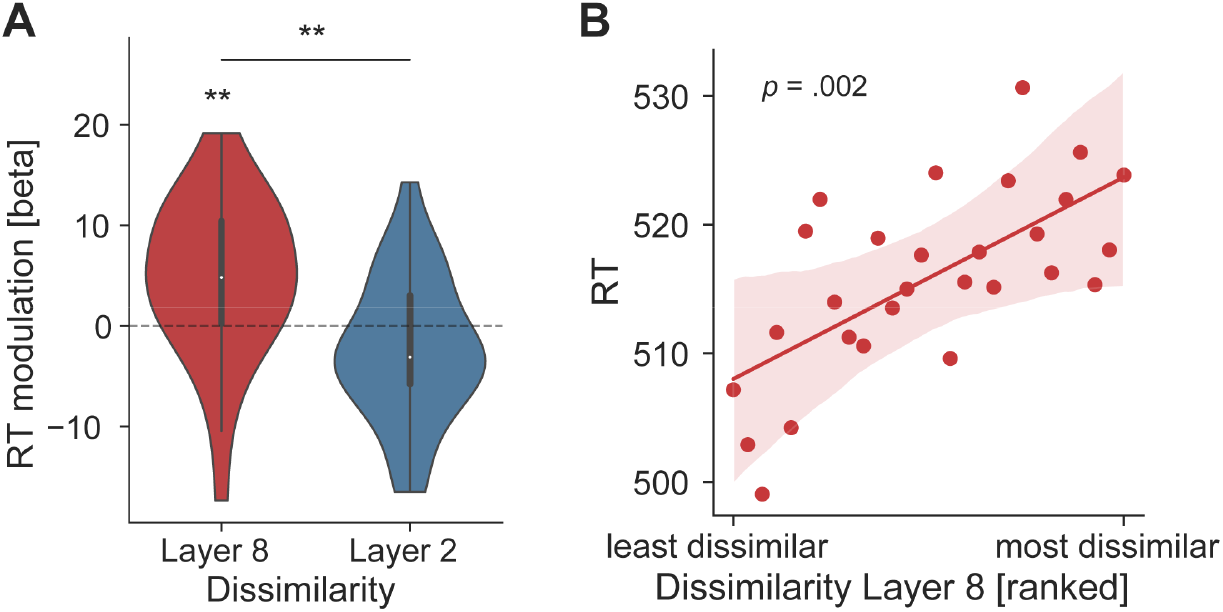
Behavioural response slowing correlates with high-level visual surprise. **A)** Behavioral responses were slower the more dissimilar surprising images were in terms of high-level, but not low-level visual features. Depicted are the slope coefficients (betas) regressing RT onto layer 8 (red) and layer 2 (blue) dissimilarity. Positive betas indicate response slowing the more dissimilar the seen unexpected image was compared to the expected stimulus. **B)** Depicts RT as a function of ranked dissimilarity (from least to most dissimilar). Dots represent the averaged RT for each dissimilarity rank. Results show a positive monotonic relationship between RT and layer 8 dissimilarity, indicating that unexpected stimuli that were more surprising in terms of high-level visual features resulted in slower behavioral responses. ** *p* < 0.01.

## Discussion

Hierarchical PP theories (Bastos et al., 2012; Clark, 2013; Friston, 2005, 2009; Rao & Ballard, 1999) have received significant attention as they propose a fundamental framework for cortical computation. Numerous studies have corroborated the main tenets of predictive processing, such as demonstrating that sensory responses to surprising inputs are enhanced compared to expected ones, likely reflecting larger sensory prediction errors (Alink et al., 2010; Kok et al., 2012; Meyer & Olson, 2011; Ramachandran et al., 2016; Richter et al., 2018). However, while evidence for this core mechanism of predictive processing has been shown across modalities, paradigms and species (de Lange et al., 2018; Heilbron & Chait, 2018; Walsh et al., 2020), it remains unknown what kind of surprise is reflected in these putative prediction errors. Here we set out to elucidate the nature of the surprise scaling of visual prediction errors and thus what information is predicted across the visual hierarchy.

### Visual prediction errors scale with high-level visual surprise

Using fMRI and representational distance measures derived from a visual DNN, our data showed that throughout multiple visual cortical areas sensory responses to unexpected images scale with the representational distance of a seen unexpected stimulus relative to the expected input. Specifically, responses monotonically scaled with high-level visual feature surprise: the larger the high-level deviation from expectation, the larger the prediction error response. Interestingly, and in contrast to feedforward processing, even early visual areas, such as V1, predominantly responded to high-level visual over low-level surprise, while being best explained in terms of low-level features in cases where no prediction was possible (localizer). These results are in line with a recent study demonstrating that firing rates in macaque V1 correlate with predictability of high-level and not low-level visual features (Uran et al., 2022). The increased prediction error to high-level visual surprise in EVC, a region not known for tuning to high-level visual features during feedforward processing, suggests that predictions are relayed top-down and hence result in the observed inheritance of feature surprise from higher areas in earlier processing stages. Thus, our results support and extend previous studies by demonstrating that (1) top-down inheritance during predictive vision generalizes across species and recording modalities, and (2) crucially appears to be a general principle of visual sensory processing evident across multiple cortical areas from EVC over LOC to the highly specialized areas in the face processing system (Issa et al., 2018; Schwiedrzik & Freiwald, 2017). Additionally, our results further demonstrate that predictive signatures and top-down inheritance of high-level feature surprise can arise, at least in humans, with little exposure to the predictive regularities, requiring only several dozen exposures rather than extensive exposure as in the case of studies in non-human primates (Schwiedrzik & Freiwald, 2017). This flexibility of the visual system to learn and rapidly utilize novel sensory priors to generate high-level predictions to inform sensory processing in earlier stages of the hierarchy further supports the hypothesis that top-down inheritance is a ubiquitous and general principle of visual processing.

What kind of mechanism may underlie the here observed modulations of sensory responses? An inherent limitation of fMRI BOLD is the temporal resolution. However, similar reports of prediction error responses in macaques (Schwiedrzik & Freiwald, 2017; Uran et al., 2022), suggest that a late stage of the neural response is modulated by top-down predictions. Albeit speculative, it is plausible that we observed the fMRI BOLD correlates of a similar late-stage top-down modulation of sensory processing. This interpretation aligns well with our observation that the feedforward response, mostly reflected during the prediction-free localizer, is dominated by local tuning properties (e.g., low-level features in V1), while recurrent processing due to prediction during the main task, relying on feedback and reflecting high-level visual surprise, takes time to arise given the necessary computations and signal relaying across multiple cortical areas. In other words, combined our results suggest that following the activation of high-level areas, during the initial feedforward sweep, a prediction is subsequently relayed down the processing hierarchy modulating the sustained phase of neural responses across earlier areas.

Elaborating further on this account, one possible explanation for the observed results is that predictions allow the visual system to settle into a valid perceptual interpretation faster and more efficiently, because predictions match the bottom-up inputs. In contrast, unexpected input will result in slower and less efficient neural processing, potentially requiring more extensive and slower recurrent processing for the visual system to derive a valid interpretation of the current inputs. The longer temporal extends but also lower efficiency of sensory inference in the case of unexpected input could then contribute to the larger BOLD signal observed here. But why would high-level instead of low-level visual feature predictability primarily modulate sensory responses? Perceptual interpretations can be constrained top-down, with higher cortical areas constraining lower areas via feedback connections (Friston, 2005). Hence, on this account, the key features for facilitating perceptual interpretation may be represented in terms of the high-level visual features encoded in the higher cortical areas sending the feedback signals. Thus, the more difficult a seen object is to reconcile with predictions by these higher-level visual areas the larger BOLD the responses (i.e., a monotonic increase), because arriving at a valid interpretation is slower and less efficient with less reliable feedback, requiring more recurrent processing across the visual hierarchy to update predictions to match the current inputs. In other words, the degree of belief updating, reflected in the prediction error magnitudes, appears to be contingent on high-level visual surprise across the visual hierarchy, including in V1.

Our analyses also demonstrated that the modulation of prediction error responses by high-level visual surprise was not explained by the task-relevant animacy category dimension or by word-level (semantic) surprise. These results thus suggest that visual prediction errors are predominantly influenced by high-level visual and not abstract linguistic or response-related surprise. This preference aligns with our earlier proposition of facilitated perceptual inference, with visual prediction errors scaling primarily with visual surprise due to the role of top-down feedback in constraining perceptual interpretations in lower areas. Consequently, because high-level regions in the ventral stream primarily encode advanced visual features, the surprise signal is expressed in terms of these visual features instead of abstract non-visual representations.

### High-level visual predictions aid perceptual inference and consequently behavioral responses

Besides sensory response modulation, high-level visual surprise was also seen to slow behavioral responses. This suggests that increased visual prediction errors may not merely reflect an epiphenomenon of predictive processing but may translate into tangible behavioral effects. Our account, that valid predictions expedite perceptual inference, could also explain why high-level visual surprise may correlate with response slowing. Essentially, slower perceptual inference could directly translate into slower behavioral responses because perceptual inference ought to conclude before response initiation. However, this interpretation is speculative as our data do not provide causal evidence, and alternative interpretations, such as slower responses due to more difficult decision making for unexpected stimuli cannot be ruled out.

That said, this account is consistent with the observation that we found reliable upregulations of prediction error amplitudes by high-level visual (layer 8) surprise in the visual system, but not in the motor system or other areas outside the visual system, such as inferior frontal gyrus or anterior insula, known to generate prediction errors especially when predictions are task-relevant (Fazeli & Büchel, 2018; Ferrari et al., 2022; Loued-Khenissi et al., 2020; Richter & de Lange, 2019). This suggests that the modulation by high-level visual surprise primarily concerns facilitated perceptual inference rather than facilitated decision making or response initiation. Yet, this does not mean that predictions do not facilitate these processes as well. Our results concern the modulation of neural responses for different unexpected inputs. The contrast of unexpected compared to expected inputs (i.e., expectation suppression) can be found in Supplementary Figure 3 and matches previous studies showing additional prediction error signatures in decision, attention and motor processing related areas (Fazeli & Büchel, 2018; Ferrari et al., 2022; Loued-Khenissi et al., 2020; Richter & de Lange, 2019), suggesting that prediction facilitates processing across multiple cortical systems.

### Flexible prediction (error) tuning

While we found no reliable upregulation of prediction error magnitudes by any of the control models (also see: Supplementary Figure 1), this does not imply that the visual system exclusively encodes high-level visual surprise. Although we dismissed that animacy category explained our results, this does not rule out that task requirements shaped the acquisition and generation of predictions, and consequently the scaling of prediction errors. Indeed, it is possible that the focus on high-level features, due to the task requirements, shaped what kind of features were predicted, given the substantial effect that tasks can have on visual processing (Harel et al., 2014). Thus, our results are consistent with recent models of adaptive efficient coding (Młynarski & Tkačik, 2022). Specifically, because our task required a high-level decision, such task demands might be reflected in the adaptive compression (silencing) of non-task relevant sensory representations due to top-down feedback, thereby resulting in the observed scaling of early visual responses with high-level visual surprise. Moreover, from an evolutionary perspective it is advantageous to develop flexible predictions, allowing the visual system to adapt to environmental requirements. In line with this hypothesis, recent evidence suggests that semantic (word level) priors can be used to generate category specific sensory predictions, even in early visual cortex (Yan et al., 2023). Therefore, more abstract, semantic representations could potentially modulate visual prediction errors under certain conditions.

On the other hand, it is also plausible that visual prediction errors can reflect low-level features, particularly if required by the environment. Indeed, in the present data small additional clusters across multiple areas scaled with intermediate-level surprise (mostly layer 3 and 4), suggesting variety in the surprise reflected in visual prediction errors. Moreover, a different task, focused on low-level features, may result in a different pattern of prediction error tuning, representing low-level surprise instead. However, it is likely that there are limits to this flexibility imposed by the architecture and representational constraints of the visual system. As argued above, high-level visual predictions may be relayed top-down and constrain visual interpretations in lower areas, because neurons in higher visual areas are tuned to high-level features. In contrast, it is less likely that high-level visual areas have the necessary representational structure and acuity to predict detailed low-level features, and hence these areas may be unable to constrain visual interpretations in EVC for low-level visual features to the same degree as for high-level predictions. Nonetheless, we believe that it is essential for future work to chart the extent to which prediction error tuning is flexibly adjusted to reflect environmental and task demands.

### Limitations

There are some limitations to the present research. A superior low-level visual feature model may have yielded prediction error modulations by such features. In addition, the specific stimulus set could have discouraged or obscured low-level surprise modulations. Nevertheless, we demonstrated that early DNN layers best explain feedforward visual activity in EVC during the prediction-free localizer runs suggesting that the employed model does reliably explain visual responses for our stimulus set. Moreover, we chose a commonly used visual DNN. This and similar models have repeatedly been shown to share representational geometry with visual cortex (Eickenberg et al., 2017; Guclu & van Gerven, 2015; Khaligh-Razavi & Kriegeskorte, 2014; Mehrer et al., 2021). Finally, there is no evidence for a strong positive, but sub-threshold, modulation of visual responses by low-level visual surprise evident in either the whole-brain or ROI results. This suggests that it is unlikely that a quantitative issue, such as low statistical power due to a subpar low-level feature model or non-ideal stimulus set for evoking low-level visual predictions, explains the current results. Nonetheless, future work could improve on the generalization of the present results by utilizing different, ideally more complex stimulus sets and improved feature models, thereby further investigating whether (other aspects of) low-level features may modulate visual prediction errors.

The word category (semantic) model has similar limitations. While word2vec and related models have been widely used to index semantic dissimilarity (Huth et al., 2016; Mitchell et al., 2008; Pereira et al., 2018), they may no longer constitute state-of-the-art models. Thus, given a different semantic feature model or task, visual prediction errors could be modulated by semantic surprise. However, while improvements in the semantic model are possible, it seems unlikely that incremental improvements in the model can fully account for the observed results, especially in EVC, given that semantic modulations tend to arise later in the processing hierarchy. In sum, while we cannot rule out that in some circumstances other models, stimuli, or tasks could result in low-level visual or linguistic semantic features modulating sensory prediction errors, our data indicates a propensity of the visual system to scale prediction errors in terms of high-level visual surprise.

### Adaptation and attention-based accounts

One concern in predictive processing studies is differentiating prediction from stimulus repetition effects such as neural adaptation. Stimulus adaptation is not a viable explanation for the present results, because all stimuli were shown equally often. Moreover, our results concern modulations of prediction error responses to unexpected stimuli based on how different they were from the expected stimuli, thereby further ruling out repetition frequency or adaptation as explanations.

Another possible interpretation is that surprising stimuli capture attention, which subsequently amplifies neural responses (Alink & Blank, 2021). While we cannot conclusively rule out this alternative account, there are multiple factors suggesting that it is not the primary factor. First, we observed the most pronounced modulation of prediction error magnitudes in EVC. If attention were the main driver, we may instead expect modulations to be more uniformly distributed across the visual hierarchy or even more pronounced in higher areas (Buffalo et al., 2010; Reynolds & Chelazzi, 2004). Second, an attention-based account must explain why attention allocation would scale primarily with high-level visual surprise, rather than semantic features or the task-relevant dimension of animacy. Even without considering these arguments, if we treat the observed modulation as an effect of attention, high-level visual surprise must be detected before attention allocation, raising the question where this surprise detection occurs if not reflected by the modulation reported here. Therefore, the predictive processing account detailed earlier appears to provide the more parsimonious explanation.

### Conclusion

We find that visual prediction errors, a key feature of hierarchical PP, primarily reflect high-level visual surprise, including in early visual cortex. These findings are consistent with predictions being relayed top-down, from higher to lower sensory areas, thereby resulting in prediction errors in early visual areas reflecting high-level visual surprise, unlike during feedforward processing. Relying on top-down feedback to constrain perception may provide computational and metabolic advantages for the sensory system, thus allowing for more efficient and rapid convergence on valid perceptual interpretations across the visual stream. Collectively, our results thereby bolster a central mechanism of hierarchical PP – the reliance of perceptual inference on prediction and prediction error generation, thus reinforcing the crucial role of predictions in perception.

## Methods

### Participants and data exclusion

In total 40 healthy, right-handed participants were recruited from the Radboud University research participation system. Of these, data from two participants were incomplete due to the participants withdrawing from the experiment. In addition, we excluded data from two further participants due to poor MRI data quality, caused by excessive motion during MRI scanning in one case and anterior coil failure in the other. Furthermore, data of three participants were excluded because of subpar behavioral performance (see *Data analysis, Data exclusion*). Thus, in total data from 33 participants (21 female, age 23.8 ± 4.5, mean ± SD) were included in the final sample.

The study followed institutional guidelines and was approved by the local ethics committee (CMO Arnhem-Nijmegen, now METC Oost-Nederland) under the blanket approval ‘Imaging Human Cognition’ (2014/288) granted to the Donders Centre for Cognitive Neuroimaging. Written informed consent was obtained before study participation and participants were compensated 10€/hour.

### Stimuli and experimental paradigm

During the experiment participants were exposed to pairs of letter cues and full-colour images of various categories while recording fMRI. On each trial the letter probabilistically predicted the identity of the image. A trial is depicted in Figure 7A. The expected image was seven times more likely to follow its associated letter cue compared to each unexpected image. The same images appeared both as expected and unexpected stimuli, with the expectation status only contingent on the cue after which the image appeared.

**Figure 7.**
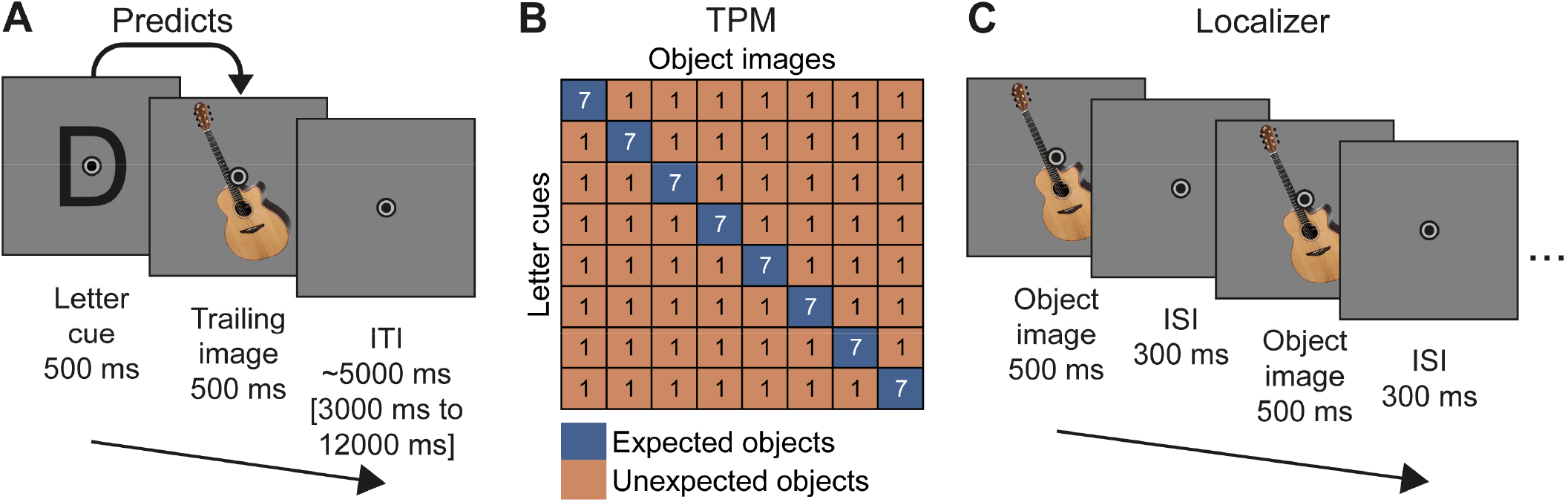
Paradigm. **A)** A single trial, showing a letter cue (500 ms) followed by an image (500 ms) and a variable ITI (∼5000 ms). The image was expected or unexpected given the preceding letter cue. Participants responded by button press to the images, indicating whether the entity in the image was animate or inanimate. **B)** Transitional probability matrix determining the associations between cues and images. Each of the eight images was associated with one of the eight letter cues. The expected image was seven times more likely to appear than any other image given its cue. Numbers in each cell indicate the number of trials per run. The specific cue-image associations were randomized and differed between participants. Moreover, the set of eight images also varied for different participants. **C)** Two cycles of a localizer trial. During the localizer one image was presented repeatedly (500 ms on, 300 ms off) for 12000 ms. The identity of the images was not predictable. Participants responded to a high brightness version of the images, which was shown once during each trial for one cycle.

#### Stimuli

Full-color images were selected from a database of 233 photographs, originally collected for a previous study (Yan et al., 2023). The image database included multiple exemplar images of various categories, including animate (dogs, dolphins, elephants, feet, hands, women and men, male and female faces, swans, tigers) and inanimate objects (cars, airplanes, churches, guitars, hammers, houses, spoons). Of the original 233 stimuli in the database 20 stimuli were excluded as outliers. Specifically, we used hierarchical agglomerative clustering to cluster image representations of layer 2 and 8 of AlexNet trained on ecoset (for more details see: *Deep Neural Network*). We then excluded any outliers, such as same category objects in a different cluster than all other exemplars of the same category in terms of layer 8 representations. From the remaining 213 stimuli eight images were selected for each participant, including four animate and four inanimate images. The selection of the eight images was optimized using the following criteria. First, we derived representational dissimilarity matrices (RDM) from the layers of interest (layer 2 and layer 8) from the DNN using correlation distance. We then randomly selected eight images (four animate and four inanimate) and calculated the variances within layer RDMs. Additionally, we calculated the across layer RDM correlation. Finally, we maximized the within layer variances and the minimized the across layer correlation. This procedure is a simple method to select image samples that maximized the detectability of effects within RDMs, while also minimizing the correlation between RDMs, thereby increasing our ability to detect distinct contributions of the RDMs from the two layers of interest. For each participant we selected a set of eight images using this procedure.

Images were presented in the center of the screen, subtending a maximum of 6 x 6 degrees of visual angle. The exact image size depended on the shape of the specific object. A fixation bulls-eye, outer circle 0.5 degrees of visual angle, was displayed on top of the center of the image.

#### Experimental paradigm and procedure

On each trial (Figure 7A) participants were presented with a letter cue for 500ms, followed by an image for 500ms without interstimulus interval. The letter cues were predictive of the image with ∼50% reliability – the transitional probability matrix (TPM) is depicted in Figure 7B. Thus, participants could predict the identity of the images given the letter. Participants were not informed about these regularities. Instead, they were tasked to categorize the entities in the images as animate or inanimate as quickly and accurately as possible. Thus, while learning the statistical regularities was not required to perform the task, the regularities could be used to facilitate task performance. To promote statistical learning, participants were required to withhold the response if the letter cue was a vowel (a, e, i, o, u), thereby directing attention also towards the letter cues. No-go letters (vowels) were not associated with any specific stimulus and appeared in addition to the regularities depicted in Figure 7B. No-go trials were discarded from all analyses. Responses on each trial were given by button press (right index or middle finger) as soon as the image appeared, with a maximum allowed reaction time of 1500ms before a trial would be considered a miss. Trials were separated by an intertrial interval of on average 5000ms (range 3000ms – 12000ms, sampled from a truncated exponential distribution), displaying only the fixation bulls-eye.

Each run, that is one continuous fMRI data acquisition, consisted of 128 trials (∼13 minutes). During a run the TPM shown in Figure 7B was presented once. Thus, each unexpected cue stimulus pair was shown exactly one time (8 x 7 combinations = 56 unexpected trials) and each expected pair was shown seven times (8 expected pairs x 7 repetitions = 56 expected trials), with the remaining 16 trials being no-go trails. Trial order was randomized, except for excluding repetitions of the same cue stimulus pair on two consecutive trials. Participants performed four runs per session and two sessions, resulting in a total of eight fMRI runs. In addition to the fMRI runs, the experiment also included an additional two behavioral blocks in each session.

The behavioral blocks were identical to the fMRI runs, except for a shorter intertrial interval (average 2500ms, range 1500ms – 7500ms) and an adjusted TPM. Specifically, expected pairs were shown three times more often during behavioral blocks compared to fMRI runs (i.e., each pair had 21 repetitions instead of 7) to facilitate statistical learning. Moreover, twice as many no-go trials were shown to compensate for the increased number of expected trials. Thus, behavioral blocks consisted of a total of 256 trials per block.

On day one participants first performed one functional localizer run (see: *Functional localizer*), followed by a short practice of the main task using different letters and images. Then, the four fMRI main task runs followed, and finally two behavioral blocks were performed outside the MRI. On the next day the order of runs was reversed. Thus, participants first did the two behavioral blocks, then the four fMRI main task runs, followed by another run of the functional localizer, and finally an anatomical scan was acquired.

#### Functional localizer

A functional localizer, depicted in Figure 7C, was performed to define object selective LOC, constrain anatomical ROI masks using independent fMRI data, and to perform RSA to validate the RDMs derived from the DNN layers of interest. The functional localizer used a block design, presenting one stimulus at a time for 12000ms, flashing every 800ms (500ms on, 300ms off), during each miniblock. Miniblock order was randomized, thus precluding prediction of the next stimulus, but excluded direct repetitions of the same stimulus. Participants were tasked to press a button whenever the image changed in brightness. The image noticeably increased in brightness (∼200%) at a random cycle exactly once per miniblock, except for during the first three and last two cycles. Each image was presented during four miniblocks per localizer run. In addition, a phase scrambled version of each image was shown, with each scrambled image being repeated in two miniblocks. As during the main fMRI task runs, images subtended 6 x 6 degrees of visual angle and a fixation bulls-eye was displayed at the center of the image throughout the entire run.

### fMRI data acquisition

MRI data was acquired on a Siemens 3T Prisma and a 3T PrismaFit scanner, using a 32-channel head coil. Functional images were acquired using a whole-brain T2*-weighted multiband-6 sequence (TR/TE = 1000/34 ms, 66 slices, voxel size 2 mm isotropic, FOV = 210 mm, 60° flip angle, A/P phase encoding direction, bandwidth = 2090 Hz/Px). Anatomical images were acquired using a T1-weighted MP-RAGE sequence (GRAPPA acceleration factor = 2, TR/TE = 2300/3.03 ms, voxel size 1 mm isotropic, 8° flip angle).

### Data analysis

#### Behavioral data analysis

Behavioral data was analyzed in terms of reaction time (RT) and accuracy. Trials with too fast (< 100ms) or too slow (>1500ms) RTs were excluded. Only trials with correct responses were analyzed for the RT analysis. RTs and accuracy were calculated for expected and unexpected image trials separately. Paired t-tests across participants on the RTs and accuracies were then performed, contrasting expected compared to unexpected images. Expectation induced behavioral benefits were calculated as RT_benefit_ = RT_unexpected_ – RT_expected_ and Accuracy_benefit_ = Accuracy_expected_ -Accuracy_unexpected_. Behavioral data was also used to reject outliers based on poor overall response accuracy and speed (for details see: *Data exclusion*).

#### Correlating RT and visual dissimilarity

We assessed the relationship between RT and visual surprise by regressing dissimilarity, as indexed by layer 8 and layer 2 of the DNN onto the RTs per participant. The obtained betas were then averaged across participants and subjected to one-sample t-tests to assess their individual effect, as well as their difference using a paired t-test. Non-parametric tests (Wilcoxon signed-rank test) were used as appropriate. For display purposes we also ranked the data for each participant into 28 dissimilarity bins (i.e., 28 unexpected image cells; see Figure 7B).

#### Effect size calculation

We provide the following estimates of effect size to support statistical inference. For t-tests we report Cohen’s *d* (Lakens, 2013), for Wilcoxon signed-rank tests matched-pairs rank-biserial correlation (*r*), and partial eta-squared 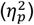 for repeated measures ANOVAs.

#### fMRI data preprocessing

fMRIprep boilerplate text: MRI data was preprocessed using *fMRIPrep* 22.1.0 (Esteban et al., 2019), which is based on *Nipype* 1.8.5 (Gorgolewski et al., 2011).

#### Anatomical data preprocessing

A total of 1 T1-weighted (T1w) images were found within the input BIDS dataset. The T1-weighted (T1w) image was corrected for intensity non-uniformity (INU) with `N4BiasFieldCorrection` (Tustison et al., 2010), distributed with ANTs 2.3.3 (Avants et al., 2008), and used as T1w-reference throughout the workflow. The T1w-reference was then skull-stripped with a *Nipype* implementation of the `antsBrainExtraction.sh` workflow (from ANTs), using OASIS30ANTs as target template. Brain tissue segmentation of cerebrospinal fluid (CSF), white-matter (WM) and gray-matter (GM) was performed on the brain-extracted T1w using `fast` (FSL 6.0.5.1; (Zhang et al., 2001) RRID:SCR_002823). Brain surfaces were reconstructed using `recon-all` (FreeSurfer 7.2.0; (Dale et al., 1999); RRID:SCR_001847), and the brain mask estimated previously was refined with a custom variation of the method to reconcile ANTs-derived and FreeSurfer-derived segmentations of the cortical gray-matter of Mindboggle (Klein et al., 2017) RRID:SCR_002438). Volume-based spatial normalization to one standard space (MNI152NLin2009cAsym) was performed through nonlinear registration with `antsRegistration` (ANTs 2.3.3), using brain-extracted versions of both T1w reference and the T1w template. The following template was selected for spatial normalization: *ICBM 152 Nonlinear Asymmetrical template version 2009c* (Fonov et al., 2009) RRID:SCR_008796; TemplateFlow ID: *MNI152NLin2009cAsym*).

#### Functional data preprocessing

For each of the 10 BOLD runs found per subject (across all tasks and sessions), the following preprocessing was performed. First, a reference volume and its skull-stripped version were generated by aligning and averaging 1 single-band references (SBRefs). Head-motion parameters with respect to the BOLD reference (transformation matrices, and six corresponding rotation and translation parameters) are estimated before any spatiotemporal filtering using `mcflirt` (FSL 6.0.5.1; Jenkinson et al., 2002). BOLD runs were slice-time corrected to 0.445s (0.5 of slice acquisition range 0s-0.89s) using `3dTshift` from AFNI ((Cox & Hyde, 1997); RRID:SCR_005927). The BOLD time-series (including slice-timing correction when applied) were resampled onto their original, native space by applying the transforms to correct for head-motion. These resampled BOLD time-series will be referred to as *preprocessed BOLD in original space*, or just *preprocessed BOLD*. The BOLD reference was then co-registered to the T1w reference using `bbregister` (FreeSurfer) which implements boundary-based registration (Greve & Fischl, 2009). Co-registration was configured with six degrees of freedom. First, a reference volume and its skull-stripped version were generated using a custom methodology of *fMRIPrep*. Several confounding time-series were calculated based on the *preprocessed BOLD*: framewise displacement (FD), DVARS and three region-wise global signals. FD was computed using two formulations following Power (absolute sum of relative motions, Power et al. (2014)) and Jenkinson (relative root mean square displacement between affines, Jenkinson et al. (2002)). FD and DVARS are calculated for each functional run, both using their implementations in *Nipype* (following the definitions by Power et al. (2014)). The three global signals are extracted within the CSF, the WM, and the whole-brain masks. Additionally, a set of physiological regressors were extracted to allow for component-based noise correction (*CompCor*, (Behzadi et al., 2007)). Principal components are estimated after high-pass filtering the *preprocessed BOLD* time-series (using a discrete cosine filter with 128s cut-off) for the two *CompCor* variants: temporal (tCompCor) and anatomical (aCompCor). tCompCor components are then calculated from the top 2% variable voxels within the brain mask. For aCompCor, three probabilistic masks (CSF, WM and combined CSF+WM) are generated in anatomical space. The implementation differs from that of Behzadi et al. in that instead of eroding the masks by 2 pixels on BOLD space, a mask of pixels that likely contain a volume fraction of GM is subtracted from the aCompCor masks. This mask is obtained by dilating a GM mask extracted from the FreeSurfer’s *aseg* segmentation, and it ensures components are not extracted from voxels containing a minimal fraction of GM. Finally, these masks are resampled into BOLD space and binarized by thresholding at 0.99 (as in the original implementation). Components are also calculated separately within the WM and CSF masks. For each CompCor decomposition, the *k* components with the largest singular values are retained, such that the retained components’ time series are sufficient to explain 50 percent of variance across the nuisance mask (CSF, WM, combined, or temporal). The remaining components are dropped from consideration. The head-motion estimates calculated in the correction step were also placed within the corresponding confounds file. The confound time series derived from head motion estimates and global signals were expanded with the inclusion of temporal derivatives and quadratic terms for each (Satterthwaite et al., 2013). Frames that exceeded a threshold of 0.5 mm FD or 1.5 standardized DVARS were annotated as motion outliers. Additional nuisance timeseries are calculated by means of principal components analysis of the signal found within a thin band (*crown*) of voxels around the edge of the brain, as proposed by (Patriat et al., 2013). The BOLD time-series were resampled into standard space, generating a *preprocessed BOLD run in MNI152NLin2009cAsym space*. First, a reference volume and its skull-stripped version were generated using a custom methodology of *fMRIPrep*. All resamplings can be performed with *a single interpolation step* by composing all the pertinent transformations (i.e., head-motion transform matrices, susceptibility distortion correction when available, and co-registrations to anatomical and output spaces). Gridded (volumetric) resamplings were performed using `antsApplyTransforms` (ANTs), configured with Lanczos interpolation to minimize the smoothing effects of other kernels. Non-gridded (surface) resamplings were performed using `mri_vol2surf` (FreeSurfer).

For more details of the pipeline, see the section corresponding to workflows in fMRIPrep’s documentation (https://fmriprep.org/en/latest/workflows.html).

#### Additional preprocessing

After preprocessing using fMRIPrep, additional fMRI data preprocessing steps were performed using FSL FEAT and Nilearn, including high-pass filtering (128s cutoff) and spatial smoothing (5mm fwhm).

#### fMRI data analysis

Univariate fMRI analyses consisted of fitting voxel-wise general linear models (GLM) to each participant’s run data, using an event-related approach. Stimuli were modelled as events of 500ms duration with the onset corresponding to the onset of the stimuli. Hence, cues, presented 500ms before stimulus onset, were not explicitly modelled. Events were convolved with a double gamma haemodynamic response function. Expected and unexpected image trials were modelled as separate regressors. Moreover, parametric modulators were added to the design matrix, reflecting how different an unexpected image was compared to the expected image. Specifically, a parametric modulator was added based on the representational dissimilarity of the unexpected compared to the expected image on a given trial in terms of layer 2 and layer 8 representations of AlexNet (see: *Deep Neural Network data* for details). Additional parametric modulators were included to serve as control variables, consisting of animacy category, word category (*word2vec*) and layer 8 distance from an untrained (random) AlexNet instance. The parametric modulators were z scored before being added to the design matrix. A regressor of no interest was included for no-go trials. First order temporal derivatives of these regressors were also added to the GLM.

Nuisance regressors were added, consisting of six standard motion parameters (rotation and translation in x, y and z), framewise displacement, CSF and white matter. All nuisance regressors were derived from fMRIPrep. To deal with temporal autocorrelation FSL’s FILM with local autocorrelation correction was used (Smith et al., 2004). Parameter estimates were averaged across runs using a fixed effects analysis, and across participants using FSL FEAT’s mixed-effects model. All fMRI analyses were performed in normalized space (*MNI152NLin2009cAsym*).

Contrasts of interest were the modulation of the BOLD response by the parametric modulators indexing the representational dissimilarity of the unexpected compared to the expected image; especially, AlexNet layer 2 and layer 8. Statistical maps were corrected for multiple comparisons using Gaussian random-field cluster thresholding, as implemented in FSL FEAT 6.0, with a cluster formation threshold of z ≥ 3.29 (i.e., *p* < 0.001, two-sided) and a cluster significance threshold of *p* < 0.05.

#### Regression of BOLD onto dissimilarity

In addition to the parametric modulation analysis, we performed a complementary analysis, regressing single trial BOLD parameter estimates (see: *Single trial parameter estimation*) onto the z scored dissimilarity metrics. This regression was performed for each participant separately, in a whole-brain and ROI fashion. The resulting slope coefficients for each voxel or ROI, indexing the modulation of BOLD responses as a function of dissimilarity were then subjected to a one-sample t-test contrasting the obtained slope against zero (no modulation). For the whole-brain analysis additional spatial smoothing of 3mm fwhm was applied (i.e., total smoothing 8mm). Finally, we colored each voxel according to which layer had the largest effect on the visual responses, indexed by explained variance. We thresholded this whole-brain analysis to a liberal z ≥ 1.96 (i.e., *p* < 0.05, two-sided) to explore the landscape of the predictive modulations.

#### Single trial parameter estimation

Single trial parameter estimates were obtained using a least squares separate approach (Mumford et al., 2012; Turner et al., 2012). A GLM was fit per trial to the BOLD data using Nilearn, where each design matrix contained a regressor for the trial of interest (iterating over all trials) and regressors of no interest for the remaining images, split by image identity, as well as a regressor for no-go trials. Nuisance regressors were also included, consisting of six motion parameters (rotation and translation in x, y and z), framewise displacement, CSF and white matter. From this we extracted the parameter estimates for each trial, with particular interest in the parameter estimates of unexpected appearances of the stimuli.

#### Region of interest (ROI) analysis

ROIs were defined a priori, based on previous studies (Richter & de Lange, 2019), and consisted of early visual cortex (V1), intermediate object-selective areas in the lateral occipital complex (LOC) and higher visual cortex (HVC) consisting of primarily of temporal occipital fusiform cortex. These three ROIs constitute well studied early, intermediate, and late ventral visual stream areas. ROI masks were defined both anatomically and functionally for each participant. First, we used Freesurfer (http://surfer.nmr.mgh.harvard.edu/, RRID:SCR_001847) for cortex segmentation and parcellation (Dale et al., 1999; Fischl, 2004), run as part of the fMRIPrep pipeline. The resulting V1 labels were transformed to native volumetric space using `mri_label2vol`. Additional atlas annotations were extracted from the Destrieux Atlas (Destrieux et al., 2010). The LOC mask was formed by combining two Freesurfer labels in the lateral occipital cortex (Middle occipital gyrus (lateral occipital gyrus), and Inferior occipital gyrus and sulcus). The HVC mask was obtained by merging three labels corresponding to higher ventral visual stream areas (Lateral occipito-temporal gyrus (fusiform gyrus), Lateral occipito-temporal sulcus, and Medial occipito-temporal sulcus (collateral sulcus) and lingual sulcus). Left and right hemisphere masks were combined into bilateral masks and dilated using a 3mm gaussian kernel. Overlapping voxels between the three ROI masks (V1, LOC, HVC) were assigned to the mask containing less voxels. We then resampled the masks to standard space (*MNI152NLin2009cAsym*) for each participant.

#### Object-selective LOC

In line with previous studies (Haushofer et al., 2008; Kourtzi & Kanwisher, 2001; Richter & de Lange, 2019), and to ensure that LOC contained stimulus selective neural populations, we further constrained the anatomical LOC masks per participant using data from an independent localizer run. In brief, we contrasted the response to intact compared to phase scrambled versions of the images, thereby obtaining voxels that respond more intact stimuli for each participant.

#### Voxel selection

In line with previous studies (Richter et al., 2018), we selected within each ROI the 200 voxels most informative about the depicted stimulus. To this end we performed a decoding analysis on the localizer data and selected the voxels affording best stimulus decoding (see: *Decoding searchlight analysis*). To ensure that our results generalize beyond the a priori selected, but arbitrary ROI size, we repeated all ROI analyses with masks ranging from 100 to 500 voxels (800 to 4000 mm^3^).

#### ROI analysis

For each subject and ROI, we extracted the contrast parameter estimates from the second level GLMs (subject level, averaged across runs) of the parametric modulators. These parametric modulators were then tested against zero (i.e., no modulation) using one-sample t-tests or Wilcoxon signed-rank tests as appropriate, as well as contrasted against one another. Resulting p-values were FDR corrected for the number of ROIs and tested parametric modulators.

#### Deep Neural Network

We used AlexNet, trained on ecoset (Mehrer et al., 2021), to derive representational dissimilarity estimates of the utilized stimuli. Early and late layers of this network have previously been shown to map well onto early and late visual cortex representations respectively (Mehrer et al., 2021). We extracted representational dissimilarities using correlation distance (1 – correlation) from all layers. RDMs of ten different instances of the DNN were averaged to minimize effects specific to any particular instance of the DNN (Mehrer et al., 2020). We were especially interested in layer 2 representing an early low-level visual feature space, and layer 8 constituting a high-level visual feature space. For more information on the DNN see: Mehrer et al. (2021). For each participant, we used the DNN derived RDMs to index how different each unexpected stimulus was compared to the expected stimulus in terms of low-level feature dissimilarity (layer 2) and high-level feature dissimilarity (layer 8). The image dissimilarity scores were z scored before being included as parametric modulators in the first level GLMs (see: *fMRI data analysis*).

#### Untrained network control analysis

In addition, we included RDMs from layer 8 of the same DNN, using the same procedure, but used an untrained (i.e., random) network instance. RDMs from this untrained network serve as a control condition to rule out DNN architecture or stimulus set specific contributions to the results.

#### Word-level and animacy category feature spaces

Additionally, we derived RDMs using word embeddings to approximate a semantic, non-visual dissimilarity metric. Specifically, we used word2vec (Mikolov, Chen, et al., 2013; Mikolov, Sutskever, et al., 2013), pre-trained on the Google News corpus (word2vec-google-news-300), to derive the pairwise dissimilarity between category words describing our image stimuli (airplane, car, church, dog, elephant, face, foot, guitar, hammer, hand, house, man, spoon, swan, tiger, woman).

#### Animacy category

As animacy was a task-relevant feature, we also derived an RDM indexing animacy. This RDM thus constitutes an object category dissimilarity and indirectly also task-response metric. To create this parametric modulator, we created a vector with zeros for unexpected stimuli with the same animacy category as the expected stimulus, and hence also response, and ones representing unexpected stimuli with a different animacy category. Dissimilarities from these control metrics were included in the first level fMRI GLM.

#### Representational similarity analysis

We validated the AlexNet derived RDMs using independent localizer data. Specifically, we used RSA (Kriegeskorte et al., 2008) to test whether the utilized DNN layer RDMs did significantly resemble the visual cortical RDMs. To this end we extracted parameter estimates for each stimulus compared to baseline from the first level GLMs of the localizer runs for each participant using a searchlight approach (6mm radius). We then z scored these parameter estimates per voxel and computed the representational dissimilarity in each searchlight sphere between the different stimuli, as indexed by the cosine distance between the vectors of the parameter estimates. This resulted in the neural RDM. We then correlated the lower triangular of the neural RDMs with the lower triangular of the DNN derived RDMs using Kendall’s Tau. For each voxel we selected the DNN layer that explained most neural variance. Finally, we also subjected the resulting correlation coefficients to one-sample t-tests across subjects for each voxel. A significant test would thus indicate that the neural RDMs and DNN derived RDMs shared representational geometry, suggesting that the DNN RDM was useful in explaining neural variance during the localizer run. We considered this a requirement for proceeding with the main task analysis of prediction error representations.

#### Decoding searchlight analysis

An additional searchlight (radius 6mm) was used to decode stimulus identity across the whole brain, using linear support vector machines (SVM). We first derived for each participant separately, single trial parameter estimate maps from the localizer run using the least squares separate procedure outlined before. On these single trial parameter estimates, using the searchlight approach, the SVM was trained and tested using 4-fold cross-validation. The labels supplied to this decoding analysis were the image identities. Thus, the resulting decoding maps indicated the ability to decode stimulus identity during the localizer runs. This decoding map was subsequently used to refine the ROI masks (see: *Region of interest (ROI) analysis*).

#### Bayesian analyses

We evaluated any non-significant frequentist tests shown in Figure 4 using equivalent Bayesian analyses to assess evidence for the absence of an effect of the low-level visual feature parametric modulator. To this end JASP 0.17.1.0 (JASP Team, 2023; RRID:SCR_015823) with default settings was used for Bayesian t-tests with a Cauchy prior width of 0.707. Qualitative interpretations of the resulting Bayes Factors were based on Lee and Wagenmakers (Lee & Wagenmakers, 2014).

#### Data exclusion

Data were excluded from analysis based on two independent criteria. First, we excluded participants due to low quality fMRI data, quantified in terms of high mean framewise displacement (FD), percentage of framewise displacement exceeding 0.2mm (FD%), high temporal derivative of variance over voxels (DVARS) and low temporal signal to noise ratio (tSNR). These four image quality metrics were derived using MRIQC (Esteban et al., 2017) for each run. For details concerning the calculation of each image quality metric see Esteban et al. (2017). Subsequently, we averaged these metrics across runs within participants and compared each participant to the sample mean. Participants with any image quality metric worse than the sample mean plus (or minus depending on the metric) 2 SD was rejected from further analysis. Two participants were excluded due to these fMRI image quality metrics.

In addition, we excluded participants based on subpar behavioural performance, indicating a lack of attention or task compliance. Using a similar approach as for the MRI quality metrics, we rejected each participant with an average behavioural response accuracy or reaction time 2 SD worse than the sample mean. Three participants were excluded for poor behavioral performance.

### Software and data availability

Python 3.7.4 (Python Software Foundation, RRID:SCR_008394) was used for data processing, analysis and visualization using the following libraries: NumPy 1.18.1 (Harris et al., 2020; van der Walt et al., 2011, RRID:SCR_008633), Pandas 1.1.4 (The pandas development team, 2020), NiLearn 0.9.1 (RRID:SCR_001362), Scikit-learn 0.24.2 (Pedregosa et al., 2011, RRID:SCR_002577), SciPy 1.5.3 (Virtanen et al., 2020, RRID:SCR_008058), Matplotlib 3.1.3 (Hunter, 2007, RRID:SCR_008624), Gensim (Rehurek & Sojka, 2011, using word2vec, RRID:SCR_014776), and Seaborn 0.11.2 (Waskom, 2021, RRID:SCR_018132). A conda environment yml file is included with the code. MRI data was preprocessed using fMRIPrep (Esteban et al., 2019) and analyzed using FSL 6.0 (Smith et al., 2004; FMRIB Software Library; Oxford, UK; www.fmrib.ox.ac.uk/fsl; RRID:SCR_002823). Whole-brain results were visualized using Slice Display (Zandbelt, 2017), a MATLAB (2022b; The MathWorks, Inc, Natick, Massachusetts, United States, RRID:SCR_001622) data visualization toolbox. Stimuli were presented with Presentation software (version 20.2, Neurobehavioral Systems Inc, Berkeley, CA, RRID:SCR_002521). All data, stimuli, as well as experiment and analysis code required to replicate the results reported in this paper will be shared upon publication in a peer-reviewed journal.

## Acknowledgements

We thank Maartje J. Graauwmans for assistance with data acquisition, Daniel Anthes with DNN programming and Karl Friston for helpful discussion of our results. This work was supported by the ERC Consolidator Grant 2020 (101000942) awarded to FPdL, as well as an ERC starting grant (TIME, Project 101039524) to TCK. Views and opinions expressed are however those of the author(s) only and do not necessarily reflect those of the European Union or the European Research Council. Neither the European Union nor the granting authority can be held responsible for them.

## Supplementary information

**Supplementary Figure 1.**
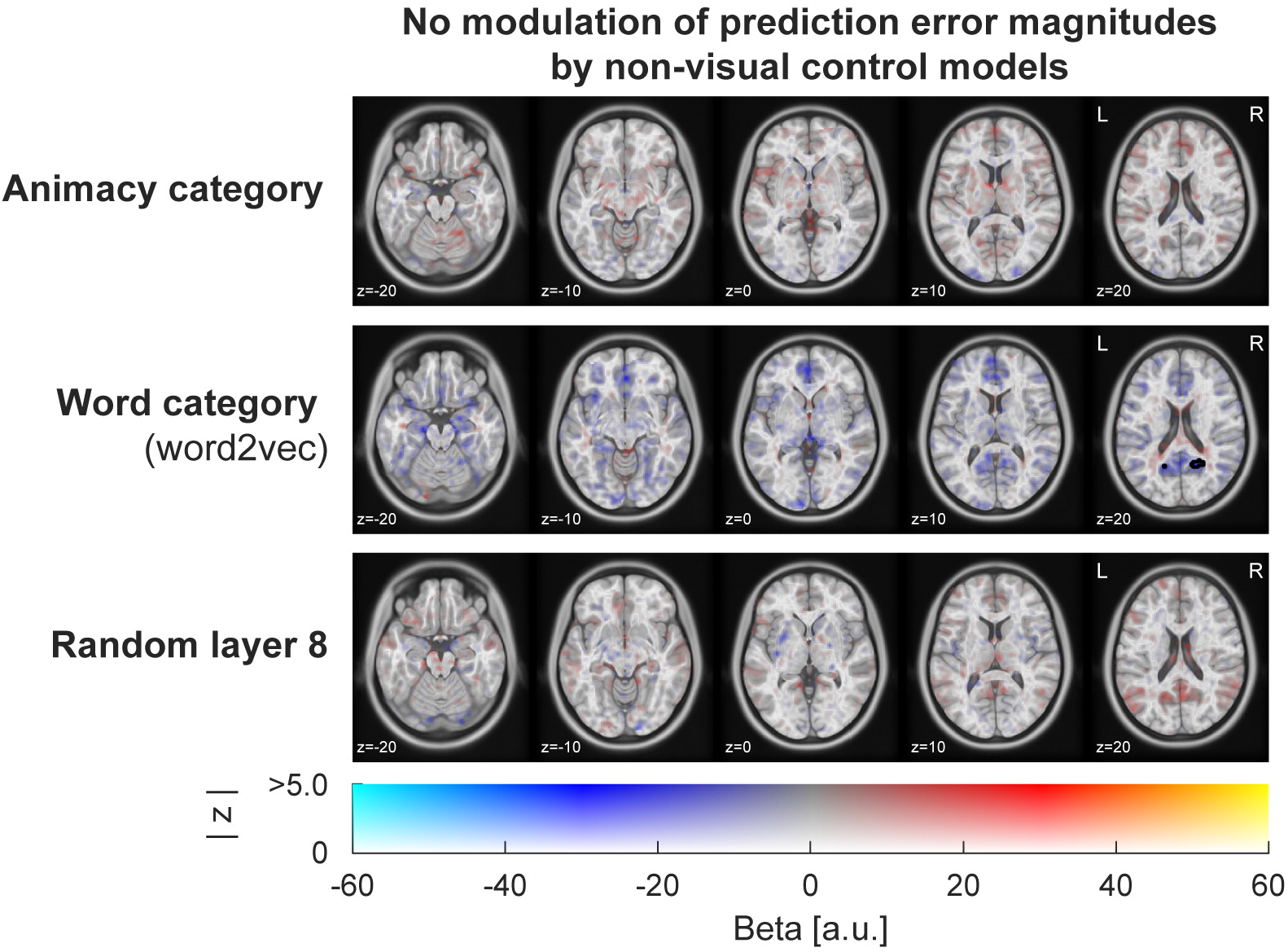
Whole-brain results assessing the modulation of surprise responses as a function of feature dissimilarity indexed by animacy category (top row), word-level semantic surprise (middle row), and a random (i.e., untrained) but otherwise identical visual DNN instance (bottom row). Results show no reliable modulation by any of these control models anywhere in cortex, except for a small negative modulation by word category surprise (word2vec) in precuneus cortex, outside of stimulus driven voxels. Thus, unexpected stimuli of a *similar* semantic category as the expected stimulus may elicit larger BOLD responses. This modulation could reflect an increased requirement for processing resources to distinguish different exemplars of the same category, albeit its small size and localization to voxels in superior parts of precuneus cortex, that were not stimulus-driven during the localizer run, makes an interpretation challenging. Colour indicates the beta parameter estimate of the parametric modulation, with red and yellow representing increased responses. Black outlines denote statistically significant clusters (GRF cluster corrected).

**Supplementary Figure 2.**
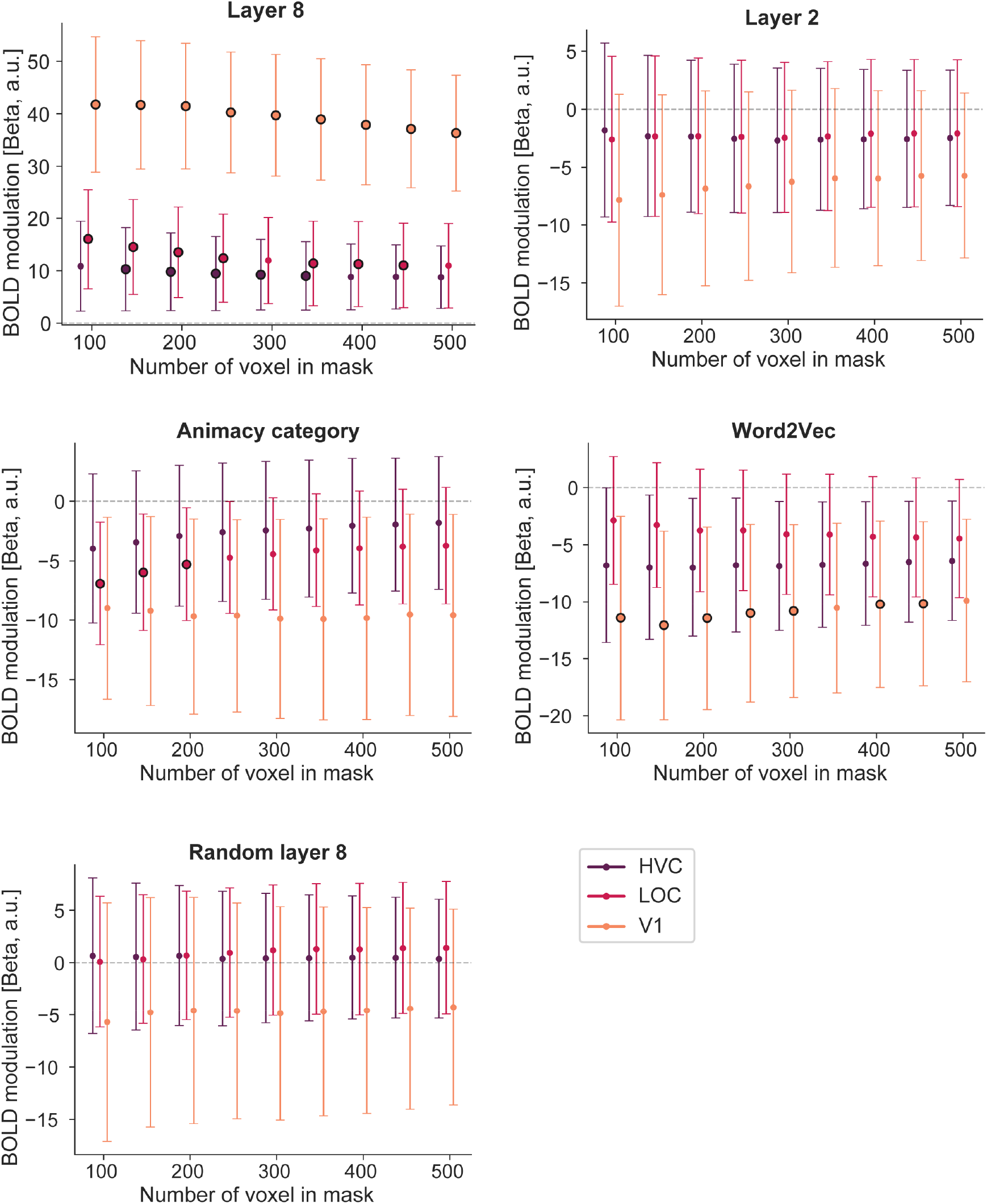
Control ROI analysis across different ROI mask sizes. Modulation of surprise as a function of high-level (layer 8; first panel), low-level (layer 2; second panel), animacy category (third panel), word level (word2vec; fourth panel), and random model (last panel) surprise. Reliable modulations of prediction error magnitudes were found for high-level visual surprise across all tested ROI sizes (100-500 voxel) in V1, as well as most mask sizes in LOC (100-250 and 350-450 voxel) and TOFC (150-350 voxel). No statistically significant modulation was found for low-level visual surprise or the random layer 8 model for any ROI or mask size. A negative modulation of BOLD responses was found for word level (word2vec) surprise in V1 for several ROI masks (150-300 and 400-450 voxel) and three small mask sizes in LOC (100-200 voxel) for animacy category. Thus, overall results closely match the results reported in Figure 5 for most mask sizes, confirming that the observed results are largely robust to variations in ROI size. Error bars depict 95% confidence intervals. Data points with black outlines indicate statistical significance at *p* < 0.05 (FDR corrected for the number of ROIs and models) compared to zero (i.e., no modulation).

**Supplementary Figure 3.**
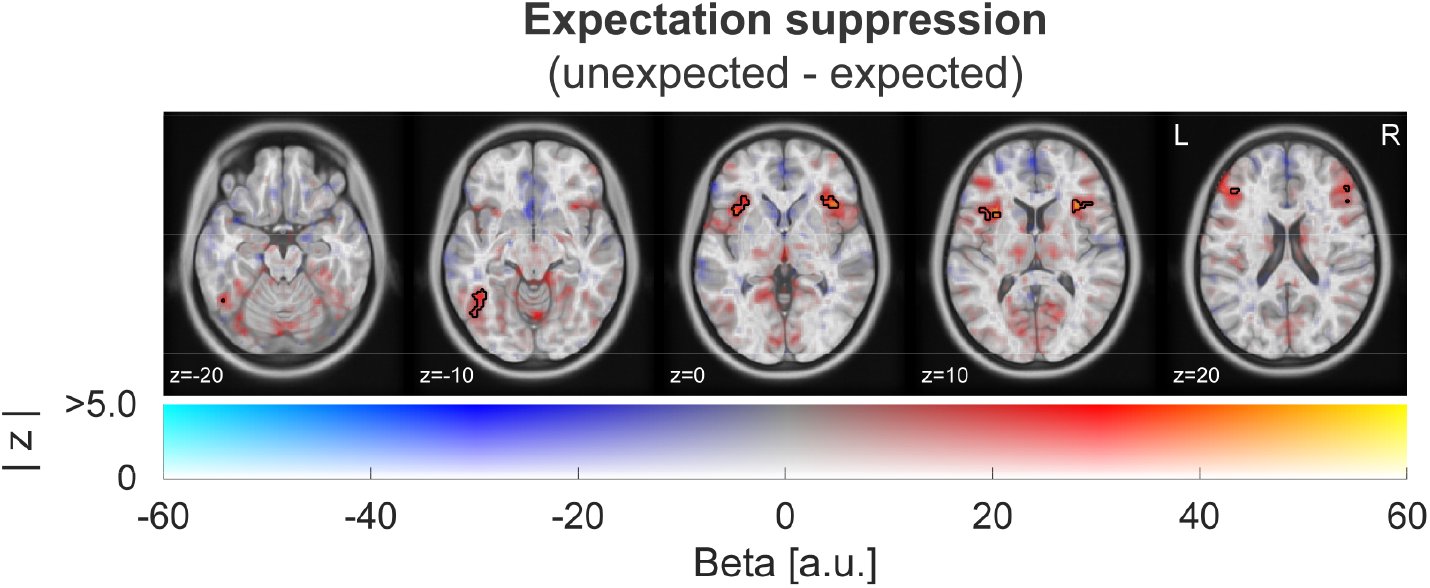
Expectation suppression. Generic prediction errors (unexpected – expected) were calculated from the voxel-wise GLM including regressors for expected and unexpected appearances of the image stimuli, as well as the parametric modulators. Here we depict the contrast unexpected – expected, thus indexing differences in neural responses contingent on whether the stimulus was expected or unexpected. Colour indicates the beta parameter estimate, with red and yellow representing increased responses to unexpected stimuli. Black outlines denote statistically significant clusters (GRF cluster corrected). Significant clusters can be seen in visual cortex, particularly in temporal occipital fusiform cortex, and outside visual cortex in anterior insula and inferior frontal gyrus. Additional significant cluster, not visible here, were found in superior parietal lobule, paracingulate gyrus, and supplementary motor cortex; see Supplementary Table 1 for details. These areas closely match previous reports of prediction error responses (Ferrari et al., 2022; Richter & de Lange, 2019).

**Supplementary Table 1.** Brain areas showing significant modulations of BOLD responses (GRF cluster corrected). Listed are the contrasts of the parametric modulators (layer 8, layer 2, word category, animacy category, and random layer 8), as well as the contrast ‘unexpected minus expected stimuli’ (Expectation suppression; Supplementary Figure 3) with corresponding area labels, numbers of voxels in the cluster, p value of the cluster, and peak z statistic. MNI coordinates indicate the X, Y, Z coordinates of the centre of gravity for the cluster, as derived by FSL FEAT’s `cluster` function, in MNI space. Area labels are based on the centre of gravity for the cluster and, especially for large clusters, additional areas encompassed by the cluster.

| Contrast | Area labels | MNI coordinates |  |  | n voxel | p cluster | max z |
| --- | --- | --- | --- | --- | --- | --- | --- |
|  |  | x | y | z |  |  |  |
| High-level visual features (layer 8) | Occipital pole; Lateral Occipital Cortex, inferior division; Occipital Fusiform Gyrus | -19 | -96 | -1 | 779 | 4.9e-19 | 5.76 |
| Low-level visual features (layer 2) | *No statistically significant clusters* |  |  |  |  |  |  |
| Word category (word2vec) | Precuneous cortex (right) | 19 | -55 | 20 | 44 | 0.0384 | -4.0 |
|  | Precuneous cortex (left) | -14 | -63 | 24 | 44 | 0.0384 | -4.2 |
| Animacy category | *No statistically significant clusters* |  |  |  |  |  |  |
| Random layer 8 | Precuneous cortex | 3 | -66 | 30 | 48 | 0.0247 | 4.0 |
| Expectation suppression (unexp. – exp.) | Superior parietal lobule; Lateral occipital cortex, superior division (left) | -36 | -56 | 47 | 839 | 9.9e-20 | 4.5 |
|  | Middle and inferior frontal gyrus; Precentral gyrus (left) | -44 | 6 | 33 | 325 | 4.0e-10 | 4.8 |
|  | Paracingulate gyrus | 1 | 13 | 48 | 265 | 9.8e-9 | 4.9 |
|  | Superior parietal lobule; Lateral occipital cortex, superior division (right) | 31 | -67 | 37 | 232 | 6.0e-8 | 5.3 |
|  | Frontal operculum cortex; Anterior insula (left) | -37 | 18 | 3 | 153 | 8.1e-6 | 4.4 |
|  | Inferior temporal gyrus; Lateral occipital cortex, inferior division | -45 | -58 | -13 | 150 | 9.8e-6 | 5.0 |
|  | Frontal operculum cortex; Anterior insula (right) | 39 | 22 | 2 | 136 | 2.5e-5 | 4.2 |
|  | Middle and inferior frontal gyrus (right) | 48 | 29 | 24 | 110 | 1.0e-4 | 4.6 |
|  | Precentral gyrus (right) | 42 | 7 | 32 | 96 | 1.0e-4 | 4.2 |
|  | Middle and inferior frontal gyrus (left) | -41 | 33 | 18 | 53 | 0.0175 | 4.2 |

**Supplementary Table 2.** Results of one sample t-tests and Wilcoxon signed rank test contrasting parameter estimates of the parametric modulators against zero (no modulation). We found reliable modulations by the high-level visual model (Layer 8) throughout all three ROIs, encompassing early (V1) and high level (HVC). P values are FDR corrected.

| ROI | Contrast | Test statistic | P value | Effect size |
| --- | --- | --- | --- | --- |
| V1 | Layer 8 | $t_{(32)} = 6.79$ | $p < 0.001$ | $d = 1.18$ |
| V1 | Layer 2 | $W = 190$ | $p = 0.199$ | $r = -0.32$ |
| V1 | Animacy category | $t_{(32)} = -2.32$ | $p = 0.068$ | $d = -0.40$ |
| V1 | Word2Vec | $W = 112$ | $p = 0.020$ | $r = -0.60$ |
| V1 | Random Layer 8 | $W = 253$ | $p = 0.719$ | $r = -0.10$ |
| LOC | Layer 8 | $W = 126$ | $p = 0.029$ | $r = 0.55$ |
| LOC | Layer 2 | $t_{(32)} = -0.68$ | $p = 0.630$ | $d = -0.12$ |
| LOC | Animacy category | $W = 137$ | $p = 0.039$ | $r = -0.51$ |
| LOC | Word2Vec | $t_{(32)} = -1.37$ | $p = 0.298$ | $d = -0.24$ |
| LOC | Random Layer 8 | $t_{(32)} = 0.21$ | $p = 0.892$ | $d = 0.04$ |
| HVC | Layer 8 | $t_{(32)} = 2.59$ | $p = 0.043$ | $d = 0.45$ |
| HVC | Layer 2 | $t_{(32)} = -0.70$ | $p = 0.666$ | $d = -0.12$ |
| HVC | Animacy category | $W = 232$ | $p = 0.579$ | $r = -0.17$ |
| HVC | Word2Vec | $t_{(32)} = -2.27$ | $p = 0.065$ | $d = -0.39$ |
| HVC | Random Layer 8 | $t_{(32)} = 0.19$ | $p = 0.852$ | $d = 0.03$ |

**Supplementary Table 3.** Results of paired t-tests and Wilcoxon signed rank test, contrasting the parameter estimates of the parametric modulators in a pair-wise fashion within each ROI. The high-level visual model (layer 8) modulated sensory responses significantly more than any of the four control models in V1, as well as some models in LOC and HVC. P values are FDR corrected.

| ROI | Contrast | Test statistic | P value | Effect size |
| --- | --- | --- | --- | --- |
| V1 | Layer 8 vs Layer 2 | $t_{(32)} = 8.05$ | $p < 0.001$ | $d = 1.40$ |
| V1 | Layer 8 vs Animacy category | $t_{(32)} = 5.71$ | $p < 0.001$ | $d = 0.99$ |
| V1 | Layer 8 vs Word2Vec | $t_{(32)} = 5.91$ | $p < 0.001$ | $d = 1.03$ |
| V1 | Layer 8 vs Random layer 8 | $t_{(32)} = 6.66$ | $p < 0.001$ | $d = 1.16$ |
| V1 | Layer 2 vs Animacy category | $t_{(32)} = 0.44$ | $p = 0.737$ | $d = 0.08$ |
| V1 | Layer 2 vs Word2Vec | $W = 218.0$ | $p = 0.440$ | $r = 0.22$ |
| V1 | Layer 2 vs Random layer 8 | $W = 216.0$ | $p = 0.440$ | $r = -0.23$ |
| V1 | Animacy category vs Word2Vec | $t_{(32)} = 0.28$ | $p = 0.810$ | $d = 0.05$ |
| V1 | Animacy category vs Random layer 8 | $t_{(32)} = -0.66$ | $p = 0.645$ | $d = -0.11$ |
| V1 | Word2Vec vs Random layer 8 | $W = 204.0$ | $p = 0.322$ | $r = -0.27$ |
| LOC | Layer 8 vs Layer 2 | $t_{(32)} = 4.24$ | $p = 0.001$ | $d = 0.74$ |
| LOC | Layer 8 vs Animacy category | $W = 108.0$ | $p = 0.010$ | $r = 0.61$ |
| LOC | Layer 8 vs Word2Vec | $W = 148.0$ | $p = 0.060$ | $r = 0.47$ |
| LOC | Layer 8 vs Random layer 8 | $W = 158.0$ | $p = 0.078$ | $r = 0.44$ |
| LOC | Layer 2 vs Animacy category | $t_{(32)} = 0.71$ | $p = 0.631$ | $d = 0.12$ |
| LOC | Layer 2 vs Word2Vec | $W = 261.0$ | $p = 0.780$ | $r = -0.07$ |
| LOC | Layer 2 vs Random layer 8 | $t_{(32)} = -0.64$ | $p = 0.607$ | $d = -0.11$ |
| LOC | Animacy category vs Word2Vec | $W = 238.0$ | $p = 0.640$ | $r = -0.15$ |
| LOC | Animacy category vs Random layer 8 | $t_{(32)} = -1.63$ | $p = 0.261$ | $d = -0.28$ |
| LOC | Word2Vec vs Random layer 8 | $t_{(32)} = -1.0$ | $p = 0.490$ | $d = -0.17$ |
| HVC | Layer 8 vs Layer 2 | $t_{(32)} = 2.85$ | $p = 0.028$ | $d = 0.50$ |
| HVC | Layer 8 vs Animacy category | $t_{(32)} = 2.32$ | $p = 0.081$ | $d = 0.40$ |
| HVC | Layer 8 vs Word2Vec | $t_{(32)} = 3.13$ | $p = 0.016$ | $d = 0.54$ |
| HVC | Layer 8 vs Random layer 8 | $t_{(32)} = 1.84$ | $p = 0.189$ | $d = 0.32$ |
| HVC | Layer 2 vs Animacy category | $t_{(32)} = 0.13$ | $p = 0.900$ | $d = 0.02$ |
| HVC | Layer 2 vs Word2Vec | $t_{(32)} = 1.03$ | $p = 0.488$ | $d = 0.18$ |
| HVC | Layer 2 vs Random layer 8 | $t_{(32)} = -0.65$ | $p = 0.628$ | $d = -0.11$ |
| HVC | Animacy category vs Word2Vec | $W = 199.0$ | $p = 0.291$ | $r = 0.29$ |
| HVC | Animacy category vs Random layer 8 | $t_{(32)} = -0.77$ | $p = 0.612$ | $d = -0.13$ |
| HVC | Word2Vec vs Random layer 8 | $t_{(32)} = -1.61$ | $p = 0.252$ | $d = -0.28$ |

